# Investigating the role of conformational heterogeneity in FUS-RRM fibrillation

**DOI:** 10.1101/2025.03.28.646059

**Authors:** Osama Aazmi, Akshit Rajendra Aswale, Leo Saju, Jeetender Chugh

**Affiliations:** Department of Biology, Indian Institute of Science Education and Research (IISER), Dr. Homi Bhabha Road, Pashan, Pune 411008; Department of Chemistry, Indian Institute of Science Education and Research, Thiruvananthapuram, Maruthamala P. O, Vithura, Kerala 695551; Department of Chemistry, Indian Institute of Science Education and Research (IISER), Dr. Homi Bhabha Road, Pashan, Pune 411008

**Keywords:** FUS, RRM, Dynamics, Conformational heterogeneity, Excited state, Amyloids

## Abstract

The Fused in Sarcoma (FUS) protein, previously implicated in neurodegenerative diseases, contains N- and C-terminal LC-rich regions, a zinc finger motif flanked by two RG-rich regions, and a single RNA-recognition motif (RRM). FUS-RRM monomers undergo amyloid-like aggregation, however, the detailed molecular insights into the fibrillation process are yet to be deciphered. Here, we investigated the conformational heterogeneity of FUS-RRM using NMR relaxation-dispersion experiments. We observed that the monomer (M) exists in a dynamic exchange with an excited state (ES), which gets perturbed by altering the pH. Although the overall fold of the FUS-RRM remains unperturbed at the lower pH, aggregation kinetics increase. The data suggests a coupling of the conformational heterogeneity to aggregation kinetics wherein a perturbation to ES probably acts as a switch that controls the fibrillation process under physiological and stress conditions. These results add to the understanding of the fibrillation process, thereby paving the way for a better understanding of the role of FUS in neurodegenerative diseases.

**Highlights:**

- FUS-RRM displays conformational heterogeneity in its folded monomeric state.
- A decrease in pH perturbs the conformational heterogeneity of the monomeric state.
- A lower pH condition accelerates the aggregation rate and also leads to a different fibril state.
- The conformational heterogeneity provides a wider target search space for potential lead compounds in structure-based drug discovery.

Graphical Abstract

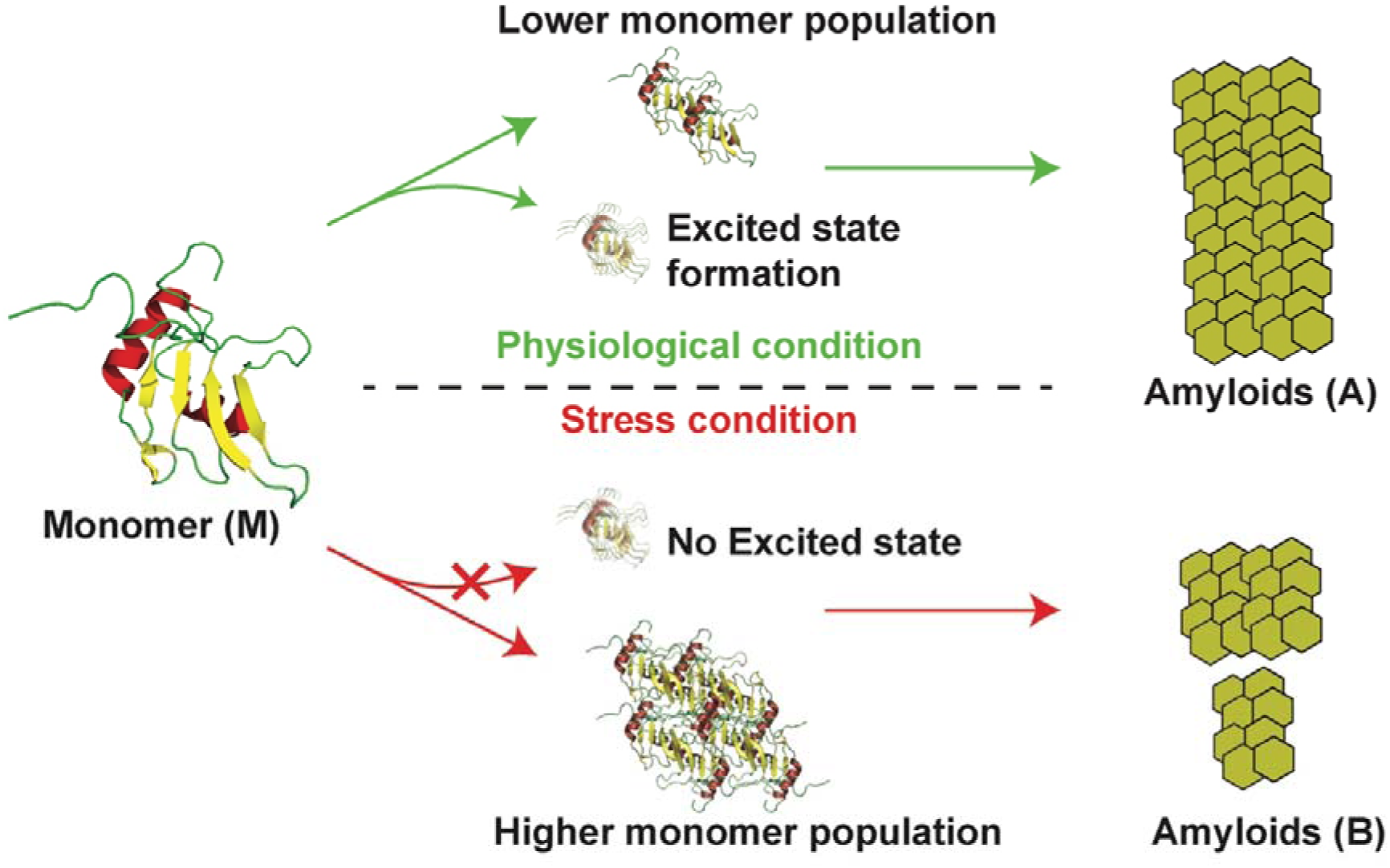

## 1. Introduction

One of the hallmarks of neurodegenerative diseases, including Alzheimer’s, Parkinson’s, amyotrophic lateral sclerosis (ALS), frontotemporal lobe degeneration (FTLD), etc., is the formation of cytoplasmic protein inclusions [1,2]. The cytoplasmic inclusions that accumulate in patients with ALS, FTLD, and Alzheimer’s disease (AD) are enriched with TDP-43 and Fused in Sarcoma (FUS) [3–8]. These RNA-binding proteins are characterized by having N- or C-terminal low complexity (LC)-rich regions and nucleic acid-binding regions known as RNA-recognition motifs (RRMs). The N- or C-terminal LC-rich regions resemble yeast prions (Sup35, Ure2, etc.) and can form amyloids [9]. The LC-rich regions are highly enriched with glutamine, asparagine, glycine, and tyrosine residues that favor liquid-liquid phase separation (LLPS) [10].

The nucleic acid-binding domain, RRM, represents the largest number of RNA-binding domains in vertebrates [11]. RRM is made up of about 90 amino acids and consists of a βαββαβ arrangement of secondary structures. The presence of ribonucleoprotein 2 (RNP2) in β_1_ and RNP1 in β_3_ (short-sequence motifs that interact with RNA) characterize RRMs [12]. Similar to LC-rich regions, the RRMs also contain amyloidogenic segments that contribute to the aggregation process associated with forming fibrillar aggregates. The nucleic-acid binding domains – TDP-43-RRM1 and 2 form aggregates under stress conditions that include pH perturbation, which tends to destabilize the soluble fraction and increase the aggregation-prone populations [13]. The TDP-43 dysfunction and aggregation have been linked through its RRM2 intermediate [14]. In a recent study, it was shown that an increase in the concentration of TDP-43 inside condensates and simultaneous exposure to oxidative stress trigger protein aggregation [15]. Similar to TDP-43, FUS-RRM has been shown to form fibril-like structures [16,17]. Thermal denaturation and renaturation studies have shown that the aggregation of FUS-RRM is irreversible [16]. However, the detailed mechanism of aggregate formation remains poorly understood for the FUS-RRM motif. We believe that understanding the protein folding pathway and the inherent conformational heterogeneity of the protein is important to obtain molecular insights into aggregate formation by FUS-RRM.

The current understanding of the protein folding mechanism – derived mainly from time-resolved FRET, single-molecule force spectroscopy in conjunction with MD simulations – indicates that the “so-called” native state of the protein is structurally heterogeneous [18,19]. Earlier studies have also shown that the folding intermediate is an ensemble of sub-populations of distinct conformations having similar free energy [20,21]. It is intriguing to understand the role of conformational heterogeneity in the misfolding and aggregation of different proteins. Indeed, computational studies have shown that the variation in the folding dynamics may influence the aggregate morphology implicated in ALS disease [22]. Previously, the role of picosecond-nanosecond (ps-ns) dynamics has been highlighted in FUS-RRM aggregation, which implies that the protein has high conformational flexibility with a low energy barrier to folded and unfolded states [16]. Insights into the energy barrier can give information about the multiple folding intermediates and their role in disease progression [21,23]. NMR is one of the best tools for detecting different protein conformations and their interconversion to multiple states. NMR relaxation dispersion experiments, including Carr-Purcell-Meiboom-Gill (CPMG), R_1ρ_, and Chemical Exchange Saturation Transfer (CEST), provide a handle to access conformational states that exist only for a short duration [24–26].

In this study, we have explored the conformational dynamics in the native state of FUS-RRM, aggregation kinetics, and pH-induced perturbation on the conformational dynamics and aggregation kinetics to understand the mechanism of fibril formation by the protein. By perturbing the exchange between the monomer population and the excited state population via pH alteration to induce stress, we show that a change in the conformational heterogeneity of the native state leads to changes in the fibril formation, thereby highlighting the role of conformational dynamics in inducing pH-dependent fibril formation of FUS-RRM. Since the study highlights the role of conformational heterogeneity of the native state in how the fibrils are formed, it suggests that the structure-based drug design approaches should not only consider the native state as a target but also include the “so-called” excited states in the target search. Thus, it opens up a new avenue for designing RRM conformational dynamics-based targets to find a therapy for neurodegenerative diseases.

## 2. Materials and Methods

### 2.1 Protein overexpression and purification

The cDNA fragment encoding FUS-RRM (aa 282-376) cloned in pET 28a vector was provided as a kind gift by Prof. Neel Sarovar Bhavesh, International Centre for Genetic Engineering and Biotechnology (ICGEB), New Delhi, India. Recombinant protein overexpression was carried out in *Escherichia coli,* as described elsewhere [27]. Briefly, the culture was induced using 0.5 mM isopropyl β-D-1-thiogalactopyranose (IPTG) at 28°C with 180 rpm for 8 hrs. The harvested culture was lysed in Tris buffer (20 mM Tris-Cl, 150 mM NaCl, and 1 mM ethylenediaminetetraacetate (EDTA), pH 8) containing lysozyme and EDTA-free protease inhibitors. Sonication of lysed cells was carried out at 60% amplitude with 5 sec ON and 5 sec OFF pulse for 3 minutes in an ice bath, cycle repeated 3 times. The cell lysate was centrifuged at 14000xg for 1 h, at 4°C, to obtain the total soluble protein (TSP). The TSP was circulated through a pre-equilibrated Ni-NTA affinity column (GE Healthcare) for 1 h and eluted using 300 mM imidazole at 4°C. Further protein was purified using a Size-Exclusion Chromatography column, Hi-Prep Sephacryl S-100 16/60 (GE Healthcare). For NMR studies, ^13^C and/or ^15^N labeled FUS-RRM samples were purified by growing bacteria in M9 media using ^13^C-glucose and ^15^NH_4_Cl as the sole source of carbon and nitrogen, respectively. The protein was concentrated in 10 mM sodium phosphate buffer, pH 6.4, containing 100 mM NaCl and 1 mM EDTA. For low pH values, the protein was buffer exchanged into 10 mM sodium acetate buffer containing 100 mM NaCl and 1 mM EDTA.

### 2.2 NMR spectroscopy

All the NMR experiments were recorded at 298 K on Ascend^TM^ Bruker AVANCE III HD 14.1 Tesla (600 MHz) NMR spectrometer equipped with a quad-channel (^1^H/^13^C/^15^N/^19^F) Cryoprobe (in-house). NMR spectra were collected with 2048×128 total points in F_2_ (^1^H) and F_1_ (^15^N) dimensions. Spectral widths of 12 ppm (^1^H) and 32 ppm (^15^N) were used, for which an acquisition time of 141 ms was obtained in the direct dimension (F_2_). All the NMR spectra were processed in NMRPipe [28] and analyzed in POKY [29]. The resonances were directly transferred from the already available assignments by Liu *et al*. and confirmed using triple resonance experiments like TOCSY, HNCA, and CBCANH [30].

Heteronuclear Adiabatic Relaxation Dispersion (HARD) experiments were recorded for different samples on the 600 MHz NMR spectrometer as described previously [31–33].

The relaxation delays used for R_1ρ_ were 0, 16, 32, 64, and 128 ms. For R_2ρ_experiments, the relaxation delays used were 0, 16, 32, and 64 ms (up to 128 ms for pH 4.6). R_1_ experiments were acquired similarly to R_1ρ_ and R_2ρ_ experiments but without using the adiabatic pulse during evolution. The delays used for the R_1_ experiment were 0, 48, 96, 192, 320, 480, and 640 ms.

R_1_/R_1ρ_/R_2ρ_ rates were calculated by fitting the peak intensities plotted against the corresponding set of relaxation delays to a mono-exponential decay model in a Mathematica script [34]. The errors were obtained using 500 Monte-Carlo simulations. R_1ρ_ and R_2ρ_ relaxation rates are dependent upon the time evolution of the effective frequency and tilt angle α,

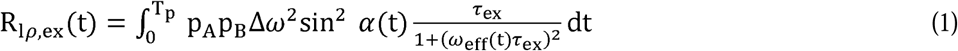

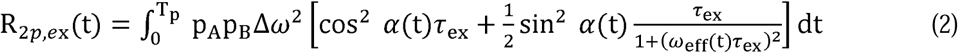

where p_A_ and p_B_ are the populations of the two exchanging sites, Δω is the chemical shift difference, and τ_ex_ is the exchange-correlation time. The R_1_, R_1ρ_, and R_2ρ_ relaxation rates obtained from the HARD experiments were used to obtain subsequent k_ex_ and p_A_ by fitting the R_1_, R_1ρ_, and R_2ρ_ data to a two-state model using numerical fittings as described previously [33].

Nuclear spin relaxation experiments (R_1_, R_2_, and [^1^H]-^15^N-nOe) were measured at the 600 MHz NMR spectrometer. The ^15^N transverse relaxation rates (R_2_) were measured using eight CPMG delays of 17, 34^∗^, 51, 68, 85, 102, 136^∗^, and 170 ms with a CPMG block length of 17 ms. The ^15^N longitudinal relaxation rates (R_1_) were measured using ten inversion recovery delays of 10, 30, 50^∗^, 100, 150, 200, 300, 450, 600*, and 800 ms. Delays marked with an asterisk were measured in duplicate to estimate errors in relaxation rates. Steady-state [^1^H]-^15^N heteronuclear nOe measurements were carried out with a ^1^H saturation time of 5 s. For the experiment without ^1^H saturation, a relaxation delay of 5 s was used. Relaxation rates were calculated by fitting the peak intensity to a mono-exponential decay against the relaxation delay in Mathematica [34]. The errors were obtained using 500 Monte-Carlo simulations. [^1^H]-^15^N nOe values were obtained as a ratio of the intensity of respective peaks of the spectra recorded with and without saturation. Errors in nOe values were obtained by propagating the errors from root-mean-square deviation (RMSD) values of baseline noise as obtained from POKY in the respective spectra [29].

All the raw relaxation data has been tabulated in supplementary tables S3 to S10.

### 2.3 Chemical shift perturbation (CSP)

CSPs between pH 6.4 and pH 4.6 were calculated in POKY [29] using the following equation:

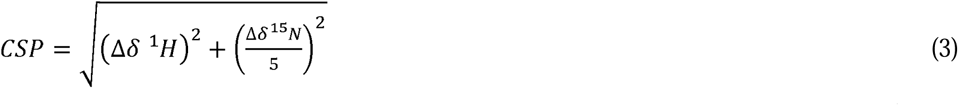

where Δδ is the chemical shift differences between the resonances in 2D ^15^N-^1^H HSQC measured at the two pH values.

### 2.4 Aggregation kinetics

pH-dependent aggregation kinetics was carried out for 200 µL reactions with different protein concentrations using 30 µM of thioflavin T (ThT) in aggregation buffer (10 mM sodium phosphate and/or sodium acetate, 100 mM NaCl, and 1 mM EDTA) at 25°C, with agitation at 60 rpm in a 96-well plate (Fluoroskan, Thermo Scientific). The fibril formation was monitored by continuously measuring the ThT intensity at 475 nm upon excitation at 440 nm. The ThT fluorescence data was measured without agitation for control experiments. The fit parameters were estimated using the least-squares method in OriginPro 8.5,

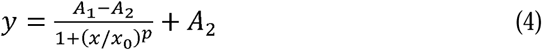

where A_1_ is the minimum, A_2_ is the maximum ThT intensity value, is 50% of the maximum ThT intensity value, and p is the power. Lag time is obtained as the derived parameter EC10 – time taken to reach the 10% of the maximum intensity attained. All the kinetics data has been reported as an average of three biological repeats (each repeat is measured from a new overexpressed and purified protein batch) and two technical repeats (each repeat is measured using the same overexpressed and purified protein) unless mentioned otherwise.

### 2.5 AFM imaging

After kinetics data were recorded, samples were centrifuged, and fibrils were resuspended in Milli-Q water. 50 µL of the sample was deposited onto a freshly cleaved mica sheet for 5 minutes, then washed multiple times with Milli-Q water and dried. AFM images were acquired in the tapping mode (Agilent technologies). Control experiments were conducted by recording AFM images on the samples before agitation. AFM data was analyzed using WSxM 5.0 software to obtain the aggregate sizes and size distribution across the AFM scan [35].

### 2.6 SEC-MALS (Size-Exclusion Chromatography with Multi-Angle Light Scattering)

SEC-MALS analysis of the purified FUS-RRM was performed on a Superdex 200 10/300 GL column (GE Healthcare) connected to an Agilent HPLC system. This system is further connected to the Wyatt Dawn HELEOS II Light scattering device with the 18-angle light scattering detector and a refractive index detector (Wyatt Optilab T-rEX). The system was calibrated with BSA at 2 mg/mL concentration. 100 µL of 1mg/ml protein sample was injected, and the molecular weight was calculated using ASTRA 2.0 software (Wyatt Technologies).

### 2.7 Circular dichroism (CD) spectroscopy

Far-UV CD spectroscopy was recorded on Jasco-815 equipped with a thermal controller for 20 µM protein in a 1 mm path length quartz cuvette with a bandwidth of 1 nm. Scans were acquired from 200-260 nm with a scanning speed of 50 nm/min. Data are represented as mean ± SD of three independent experiments after blank subtraction. The secondary structures from the CD spectrum were deconvoluted using BeStSel (Table S2) [36].

### 2.8 Fluorescence spectroscopy

Tryptophan fluorescence for different pH values with 20 µM protein was carried out in a Fluoromax-4 spectrofluorometer (Horiba Scientific). The fluorescence spectrum for each sample was obtained by exciting tryptophan at 280 nm and collecting emission spectrum from 290-500 nm with slit widths: excitation at 5 nm and emission at 1 nm. Data are represented as mean ± SD of three independent experiments after blank subtraction.

## 3. Results

### 3.1 FUS-RRM purification, characterization, and fibril formation at physiological conditions

FUS-RRM was purified with high solubility (20 mg/ml) at pH 6.4 under non-denaturing conditions, Fig. 1A. Protein characterization using SEC-MALS shows that the protein is monomeric with a molecular weight of 11.6 kDa, Fig. 1B. However, upon agitation, the monomer undergoes fibrillation in a concentration-dependent manner, Fig. 1C and S1, with an increase in the ThT intensity and a decrease in the lag time (Table S1). The increase in the ThT fluorescence intensity indicates the formation of amyloid fibrils in an otherwise folded protein [17]. The transition from folded monomer (M) to amyloid fibrils (A) was confirmed using AFM, Fig. 1D and S2. Considering only isolated fibrils whose ends can be easily detected, AFM measurements indicate the length of a fiber is 0.6 µm and a height of about 12 nm. For the protein samples without agitation, AFM measurements did not detect any fibrils (Fig. S2). The chemical shift dispersion in the ^1^H dimension of the 2D ^15^N-^1^H HSQC spectrum, marked with the resonance assignment, shows the protein is in the folded conformation (Fig. 1E).

**Figure 1.**
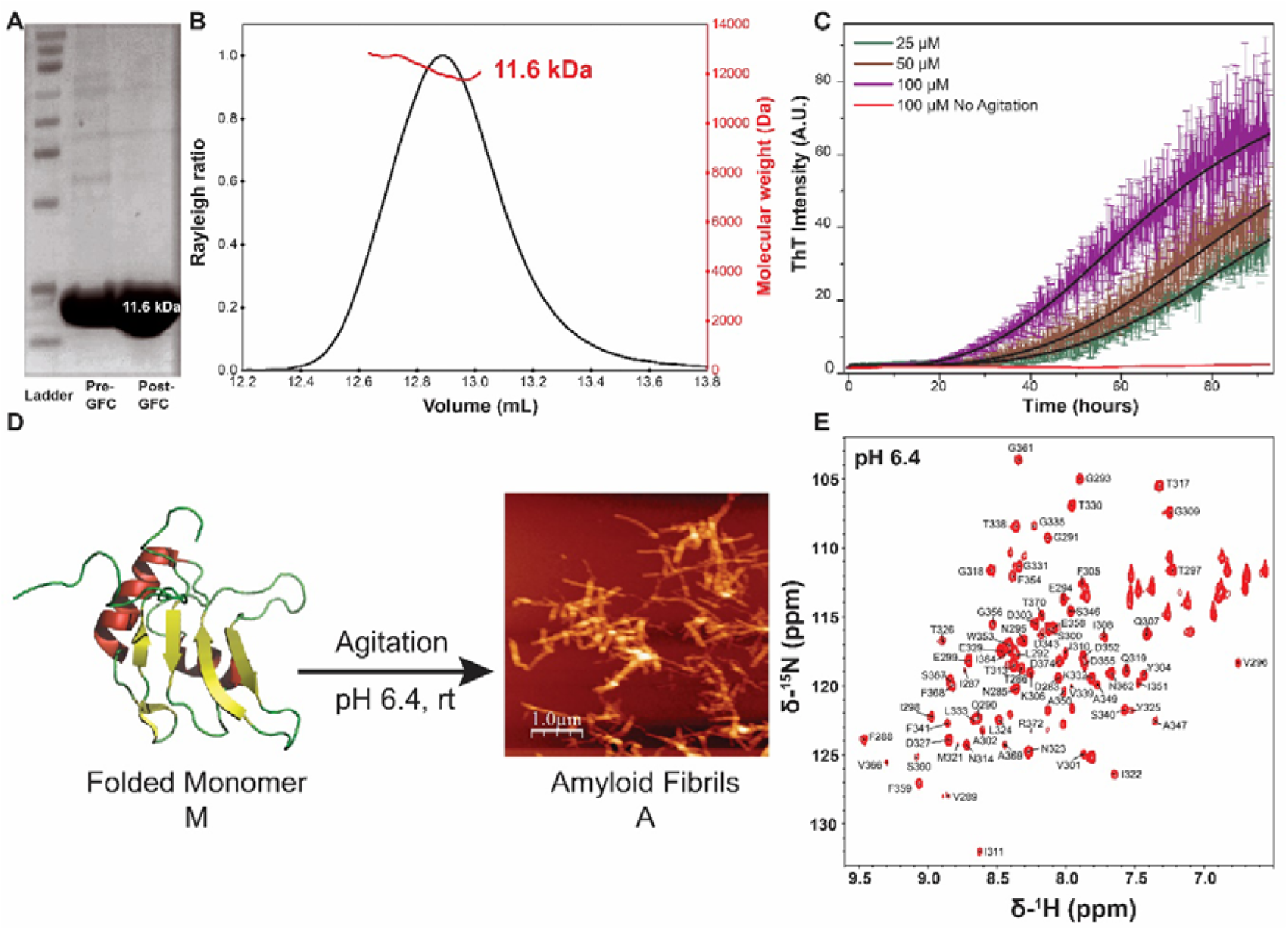
FUS-RRM characterization and fibril formation. **(A)** SDS-PAGE of purified protein. **(B)** Protein molecular weight characterization using SEC-MALS at pH 6.4. **(C)** Aggregation kinetics at pH 6.4 monitored by ThT intensity while agitating the protein at 60 rpm and 25°C, a solid line represents the fit to the kinetics data to eq (4). Data are represented as mean ± SD of three biological repeats with two technical repeats. ThT intensity at pH 6.4 does not increase when the protein is not agitated. **(D)** The folded monomer, M, undergoes fibril formation to state A upon agitation at room temperature. **(E)** Resonance assignment marked on 2D ^15^N-^1^H HSQC spectrum of RRM measured at pH 6.4.

### 3.2 The µs-ms timescale conformational dynamics

Previously, FUS-RRM has been shown to undergo irreversible unfolding and self-assembly into amyloid fibrils [16]. ^15^N backbone relaxation data (T_1_, T_2_, and heteronuclear nOe) show FUS-RRM has high conformational flexibility on ps-ns timescale. This indicates that the barrier to separate the folded and unfolded states of the FUS-RRM is relatively low, and these states exchange on a timescale faster than the CPMG timescale (k_ex_ ∼ 1000-7000 s^-1^) [16]. Here, heteronuclear R_1ρ_ (longitudinal relaxation rates in a spin-lock field) and R_2ρ_ (transverse relaxation rates in a spin-lock field) NMR relaxation dispersion experiments are measured to capture the µs-ms dynamics (faster than the CPMG timescale) [31]. Heteronuclear adiabatic R_1ρ_ and R_2ρ_ NMR relaxation dispersion (HARD) experiments are useful for detecting conformational or chemical exchange processes occurring on the fast μs-ms timescale (k_ex_ ∼ 10^4^-10^5^ s^-1^). The sensitivity of this experiment to the motion timescale can be adjusted by varying the maximum spin-lock field strength () in the adiabatic pulses for both R_1ρ_ and R_2ρ_ [32]. An adiabatic full passage (AFP) pulse is used to induce a dispersion in the relaxation rates, which are then analyzed using different models to determine the exchange rate (k_ex_) and population of alternative conformation data [31]. Since the R_1ρ_ and R_2ρ_ have differential sensitivities to the exchange process and are analyzed together with longitudinal relaxation rates (R_1_), the reliability in the fit parameters improves, much like using data measured at two magnetic fields. We have previously used these experiments to study alternative conformations in double-stranded RNA-binding proteins [33,37].

Fig. 2 shows the measured relaxation rates plotted against the residue numbers for multiple stretching factors (HS_n_) of the adiabatic pulse used to create a spinlock. For R_1ρ_, an increase in the rates with an increase in HS_n_ (Fig. 2A), and for R_2ρ_ rates, a decrease was observed (Fig. 2B). For a system that is rigid on the timescale sensitive to these experiments, no change in R_1ρ_ and R_2ρ_ rates is expected, which is observed at the terminal (D283 and D374) and a few loop residues (T317, G319) serving as an internal control. Different parameters which could be extracted using a two-state model after numerical fitting to relaxation data include k_ex_, p_M_ (population of the monomer), and delta omega Δω (chemical shift difference between the ground state and excited state), as explained in the methods section. The k_ex_ rates vary from 0.1-130 x 10^3^ s^-1^, and most residues are found to exhibit fast exchange dynamics at this timescale. Measured k_ex_ rates have been mapped onto the structure of the FUS-RRM in Fig. 2C. The population of the monomer is shown in a scatter plot, Fig. 2D. The monomer represents about 68-99% of the population in a residue-wise plot with an average value of 93.7%. Residues showing a population of excited state (ES) > 5% have been mapped on the tertiary structure of the protein (Fig. S3) to highlight the clustering of such residues on one face of the protein molecule.

**Figure 2.**
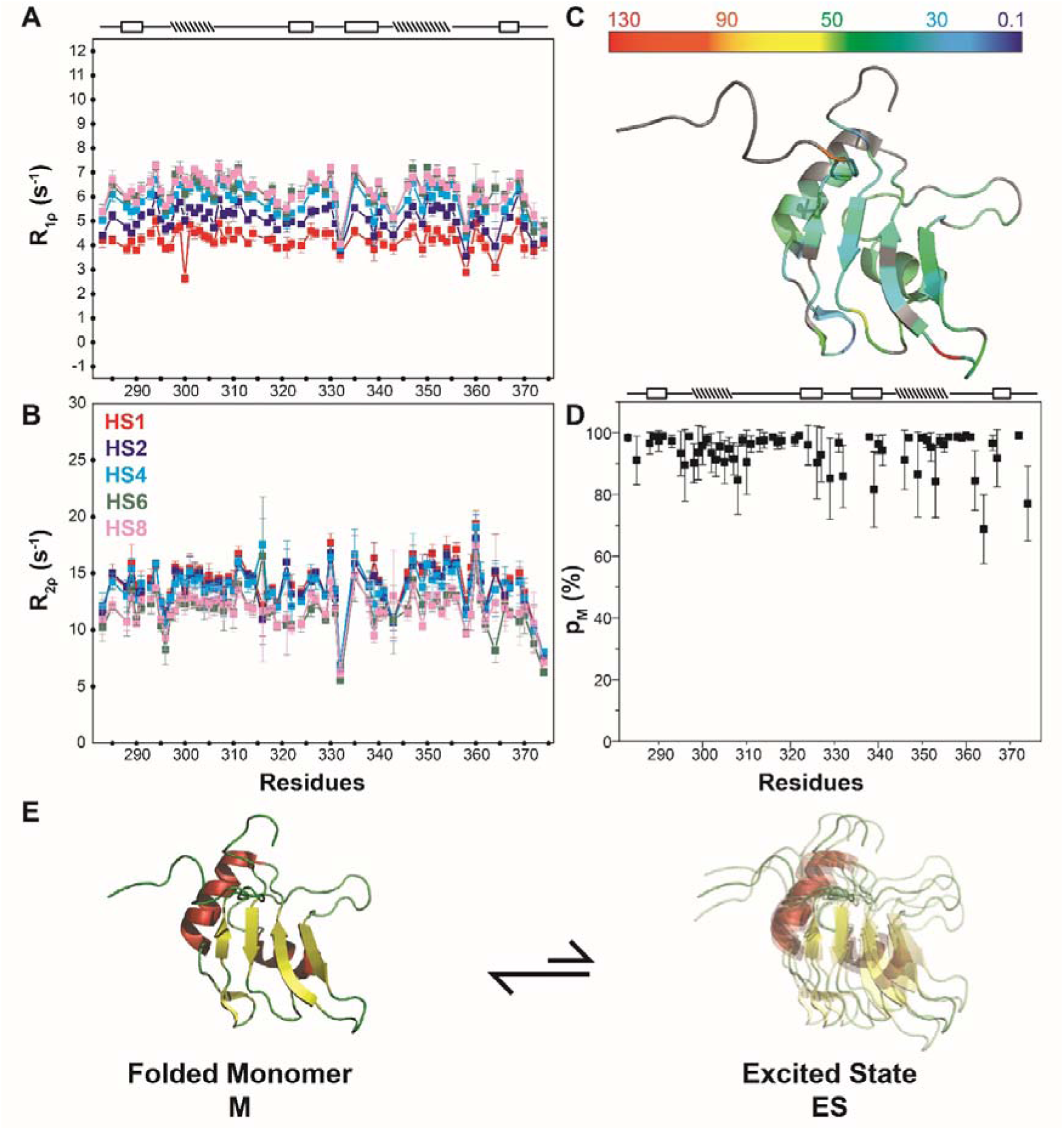
The µs-ms dynamics measured at pH 6.4. **(A)** R_1_ρ and R_2_ρ relaxation rates at 25 °C. The trends in the relaxation rates for R_1_ρ are HS1 (red) < HS2 (blue) < HS4 (sky blue) < HS6 (green) < HS8 (pink) and, **(B)** for R_2_ρ trends are HS1 > HS2 > HS4 > HS6 > HS8. **(C)** k_ex_ rates mapped to the FUS-RRM tertiary structure. The residues highlighted in grey do not fit any model and, hence, are not included in the discussion. **(D)** The population of the monomer. The secondary structure of FUS-RRM is indicated on the top of **A** and **D**, with boxed and dashed rectangles denoting strands and helices, respectively. **(E)** A schematic of an exchange between the monomer, M, and an excited state, ES.

The above results show the FUS-RRM monomer (M) exists in fast exchange with another state – the excited state (ES) (Fig. 2E). ThT kinetics and AFM data show that the M goes to the A-state upon agitation (Fig. 1D, S1, and S2). Thus, we propose that the conformational heterogeneity observed in the M-state is linked with the formation of the fibrils.

To validate this hypothesis, a perturbation of exchange between M and ES and studying its effect on the kinetics of M to A seemed a plausible strategy. If external stress perturbs the exchange dynamics between M and ES, which leads to either the state change of fibrils or the kinetics of fibril formation, it will point towards a coupling of conformational heterogeneity with the fibril formation. However, if the exchange dynamics is perturbed, and no change in the fibril formation is observed, it would suggest no such coupling exists.

We next wanted to identify a suitable perturbation that would only affect the exchange dynamics between M and ES without perturbing the overall fold of the protein. It should be noted here that since the relaxation dispersion experiment has a limited window of sensitivity, it is very much possible that there are other conformations present in the equilibrium that go undetected in this study. To measure other such conformations is not trivial and would depend on a lot of parameters, e.g., lifetime, stability, exchange rates, etc., and other biophysical experiments must be carried out for their detection. The thermodynamic stability of protein conformations results from a delicate balance between electrostatic interactions, hydrophobic forces, and the environmental cues perturbing these interactions. A change in pH is one such perturbation that can disrupt this balance, leading to structural transitions and functional alterations in proteins. Thus, we tested different stress conditions to perturb the exchange dynamics between M and ES. Out of different stress conditions (pH and ATP-binding) tried to suppress the exchange dynamics between the folded M and the ES, we observed pH 4.6 was minimally perturbing to the backbone fold and thermal stability, and effectively suppressed the exchange between the M and the ES. Thus, we used the pH 4.6 condition as an external perturbation to test our hypothesis.

### 3.3 pH-induced dynamics perturbations measured by R_1ρ_ and R_2ρ_

The adiabatic R_1ρ_ and R_2ρ_ relaxation rates were measured as a function of decreasing pH (from pH 6.4 to 4.6), shown in Fig. 3. While the dispersion in the R_1ρ_ relaxation rates (ΔR_1ρ_) did not change significantly, the one in R_2ρ_ relaxation rates (ΔR_2ρ_) decreased significantly from pH 6.4 to pH 4.6 (Fig. 2A, 3A, and S4), suggesting the reduction in the dynamics of the protein at the timescale measured by these experiments. After fitting relaxation data, k_ex_ rates and excited state population were determined. The protein indeed is rigidified at pH 4.6 at the timescale measured by adiabatic R_1ρ_ and R_2ρ_ experiments, as observed by a significant decrease in k_ex_ (0.1-2 x 10^3^ s^-1^), Fig. 3B. Similarly, the population of the ground state (M) also increased significantly (average p_M_ = 98%) with a decrease in the pH, Fig. 3C. This indicates that pH perturbation significantly quenches the conformational exchange to the ES significantly.

**Figure 3.**
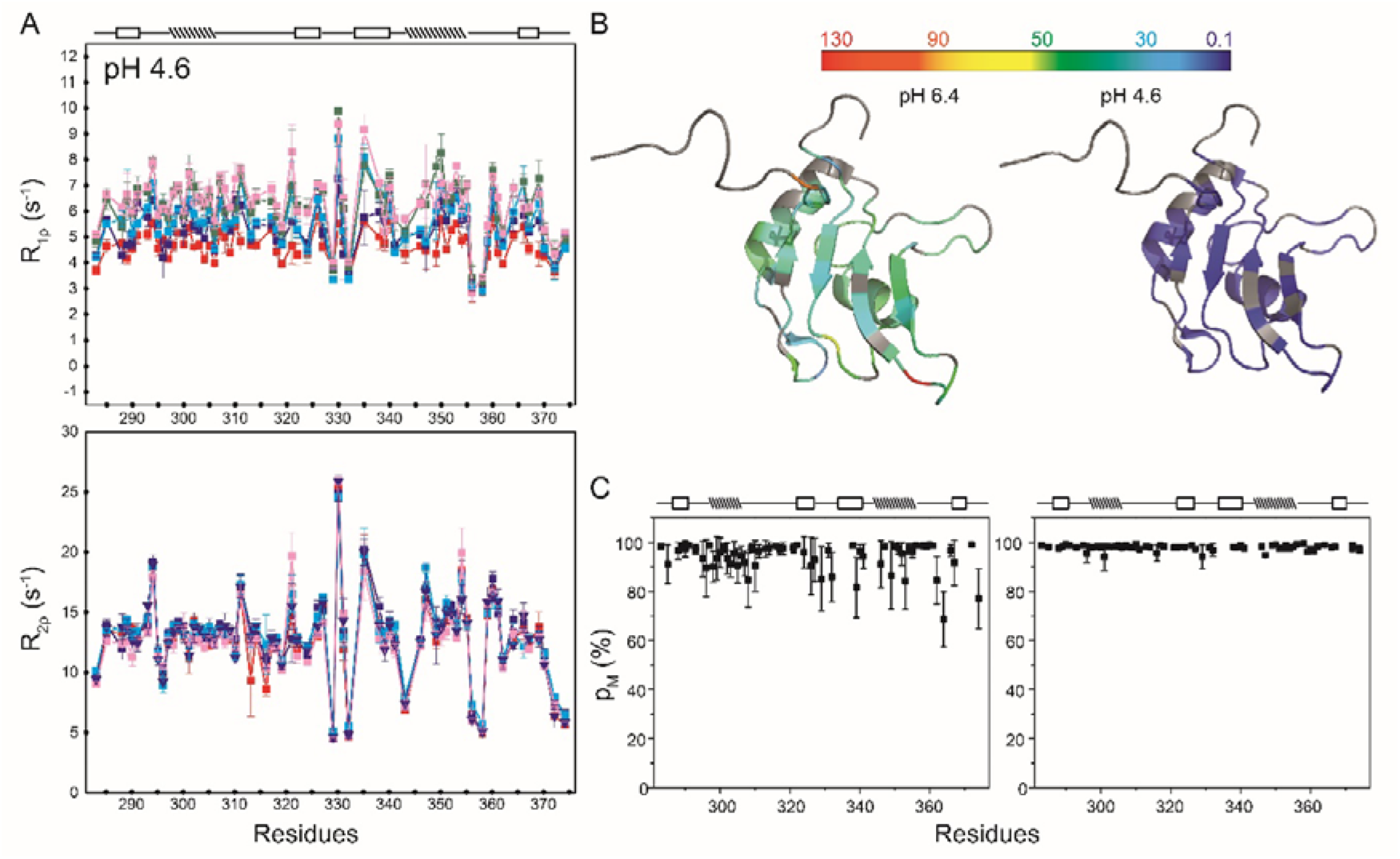
pH-induced dynamic perturbations. **(A)** R_1_ρ and R_2_ρ relaxation rates as measured for FUS-RRM monomer as a function of decreasing pH (pH 6.4 to pH 4.6) at 25 °C. **(B)** A comparison of k_ex_ rates mapped to the FUS-RRM tertiary structure for pH 6.4 and pH 4.6. The residues highlighted in grey do not fit to any model and hence are not included in the discussion. **(C)** Population of the ground state for pH 6.4 and pH 4.6. The secondary structure of FUS-RRM is indicated above the plots, with boxed and dashed rectangles denoting strands and helices, respectively.

### 3.4 pH-induced dynamics perturbations measured by R_1_, R_2_, and [^1^H]-^15^N nOe

We tested the pH-induced perturbations to the fast dynamics (ps-ns timescale motions) using traditional R_1_, R_2_, and [^1^H]-^15^N nOe experiments (Fig. S5). It can be seen that the fast dynamics – to which R_1_, R_2_, and [^1^H]-^15^N nOe experiments are sensitive – is not significantly perturbed by the pH. The average values for R_1_ remain well within one standard deviation. For the R_2_ and nOe, the average values remain within two standard deviations due to two residues in the loop being significantly perturbed. In addition to the motion of the NH bond vector, the fast dynamics also provide information on the rotational correlation time of the protein, which in turn reports on the overall size of the molecule. Thus, similar average values of R_1_, R_2_, and [^1^H]-^15^N nOe parameters at two pH values suggest that the change in the pH does not perturb the overall size of the molecule in solution.

### 3.5 pH-induced structural perturbations

For proteins like FUS-RRM, which form amyloid fibrils, changes in pH can either increase or decrease fibril formation [13]. pH-perturbations may change the secondary or tertiary structure of the protein, thereby affecting residues involved in fibril formation. CD and fluorescence spectroscopy were performed to find changes in secondary or tertiary structure. Fig. 4A shows the far-UV CD spectroscopy at pH 6.4 and pH 4.6. For both the pH values, the overall secondary structure remains largely similar (Table S2), with a prominent dip at 207 nm. Fig. 4B shows the tryptophan fluorescence for both pH values and the fluorescence maxima observed at 355 nm. This indicates that the protein backbone fold remains similar in the two pH conditions studied. Thermal stability measurements indicate a similar T_m_ of 50-55°C for a large perturbation in the pH (Fig. S6). Further pH-dependent 2D ^15^N-^1^H HSQC showed a similar protein fold, Fig. 4C. The chemical shift perturbations (CSPs) from pH 6.4 to pH 4.6 for the backbone amides were observed to be of the order of <0.5 ppm, suggesting an overall small change in the folded monomer M (Fig. 4D). The twelve residues showing CSPs above the plus one standard deviation (SD) line are considered significantly perturbed and are mapped on the tertiary structure of the protein (Fig. 4E). It indicates that out of the twelve residues, ten residues are surface-exposed and thus may not perturb the overall tertiary fold of the molecule. These results indicate that the protein structure of the state M is only minimally perturbed, while the exchange dynamics between M and ES is significantly quenched.

**Figure 4.**
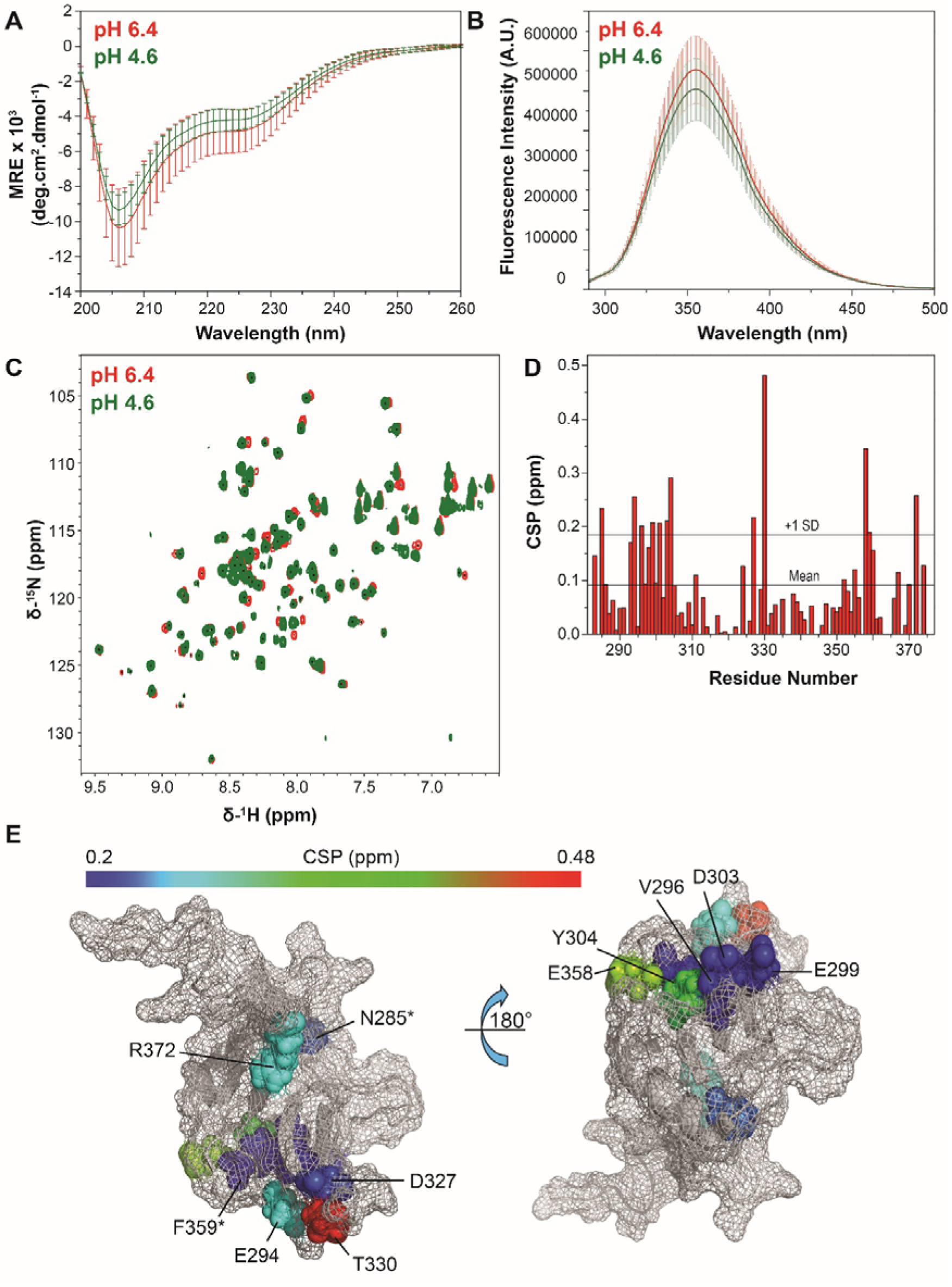
pH-induced structural perturbations in FUS-RRM. **(A)** Far-UV CD spectroscopy for pH 6.4 and pH 4.6. Data are represented as mean ± SD of three independent experiments after blank subtraction. **(B)** Tryptophan fluorescence spectroscopy for pH 6.4 and pH 4.6. Data are represented as mean ± SD of three independent experiments after blank subtraction. **(C)** An overlay of 2D ^15^N-^1^H HSQC for FUS-RRM recorded at pH 6.4 and at pH 4.6, and **(D)** Chemical shift perturbation obtained from ^15^N-^1^H chemical shifts plotted against the residue number for FUS-RRM, as described in Methods. Two horizontal lines show the mean, and the mean + standard deviation (SD). Residues showing CSPs above the SD line are considered significantly perturbed. **(E)** The twelve significantly perturbed residues obtained from (D) have been mapped on the tertiary structure to indicate that ten of these residues are surface-exposed and do not perturb the overall tertiary fold of the molecule. The color gradient shows the extent of CSPs within the 12 residues. The residues marked with stars (N285 and F359) are buried inside.

### 3.6 pH-induced perturbation on aggregation kinetics

Subsequently, we investigated the impact of pH variation on aggregation kinetics. Figs. 5A and 5B illustrate the concentration-dependent aggregation kinetics using ThT at pH 6.4 and 4.6, respectively. Notably, pH 4.6 exhibited accelerated aggregation kinetics, characterized by a significant reduction in lag time compared to pH 6.4. Specifically, the lag time decreased from 54 hours to 8.5 hours at a concentration of 25 µM (Table S1). However, there was an overall decrease in ThT intensity at pH 4.6, which could be attributed to structural or morphological differences observed in the fibrils (see below). Intriguingly, at pH 4.6, the lag time was eliminated across all concentrations.

**Figure 5.**
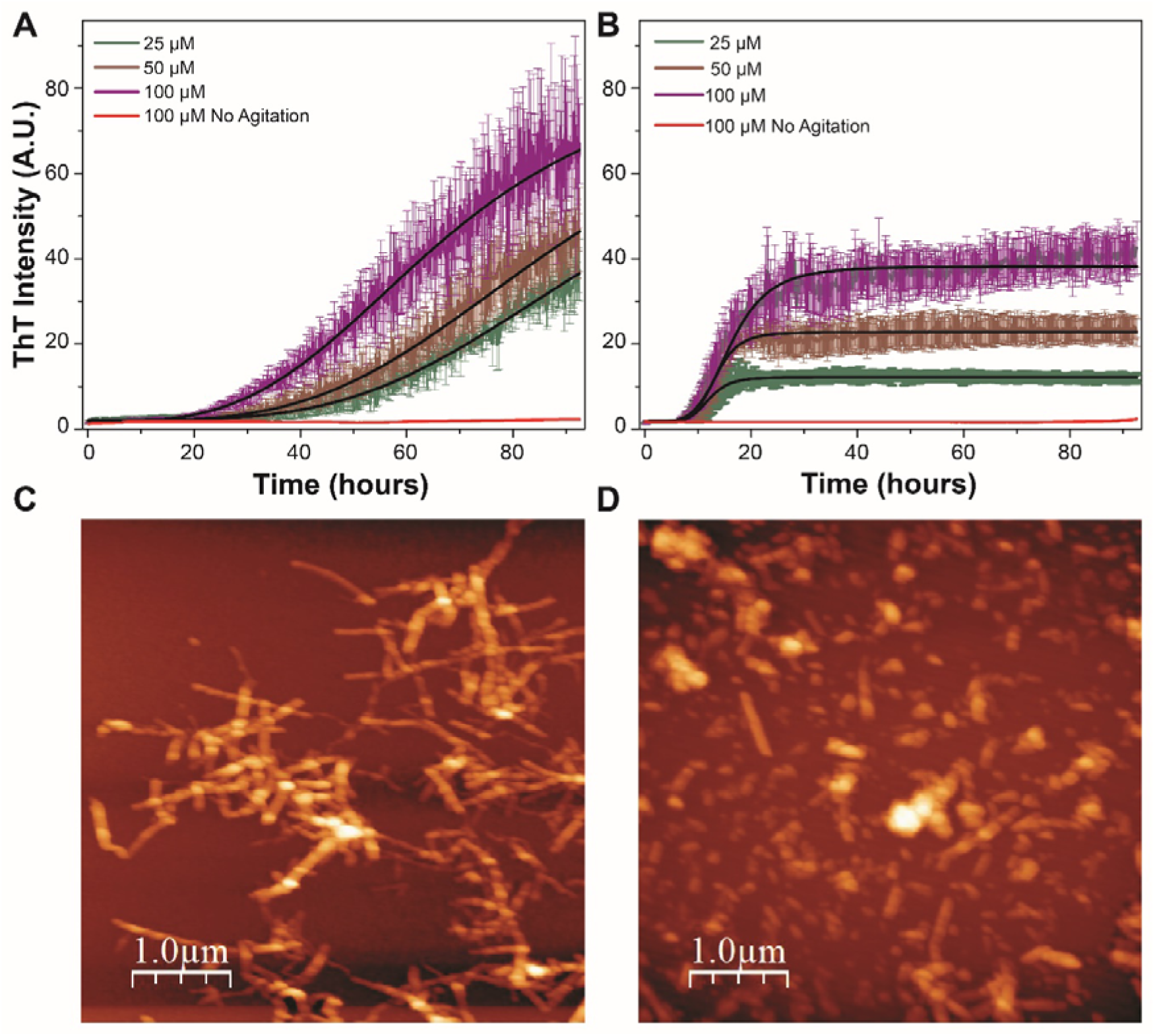
Aggregation kinetics for FUS-RRM as measured by monitoring ThT intensity while agitation at **(A)** pH 6.4, and **(B)** pH 4.6. The solid line represents the fit to the kinetics data. Fibril formation for FUS-RRM as imaged by AFM at **(C)** pH 6.4, and **(D)** pH 4.6.

### 3.7 pH-induced perturbation in the fibril state

The corresponding AFM images at the two pH values reveal observable differences in fibril morphologies. Although there are fibers formed of similar length at the lower pH (0.8 μm with a similar height as that detected at pH 6.4, AFM analysis also showed the formation of smaller fibrils and amorphous aggregates at pH 4.6. (Fig. 5C, 5D, and S2). Since the fibril morphologies appear significantly different at the two pH values, we called fibrils at pH 6.4 aggregate state A and that at pH 4.6 as aggregate state B. Similar analyses for samples without agitation do not show any fibril formation (Fig. S2). Differences in dynamics due to pH perturbations can lead to varied local folding, which may affect aggregate formation. It is tempting to speculate that the observed variability in the aggregate formation may be due to differences in the excited state structural propensities.

## 4. Discussion

In light of the intrinsic conformational exchange, aggregation kinetics, and AFM data (Fig. 5) measured at pH 4.6, we can now revisit the hypothesis. In this study, we have shown that the structure of the monomer is minimally perturbed (shown by far-UV CD, tryptophan fluorescence, 2D ^15^N-^1^H HSQC NMR spectroscopy, and ps-ns timescale motions measured by R_1_, R_2_, and [^1^H]-^15^N nOe experiments). Since perturbation in all these experiments is within the error bar, it is safe to believe that the monomer conformation is minimally perturbed. Despite the monomer conformation being similar at the 2 <u>pH</u> <u>conditions used</u>, the accelerated kinetics at pH 4.6 can be attributed to charge imbalance by the surface residues, which is highlighted by the CSP of such residues and will need further investigation. The study highlights the role of surface residues and conformational dynamics for FUS-RRM aggregation.

Monomer population (and hence effective concentration) is perturbed at lower pH due to the quench of the conformational exchange between the monomer and the excited state (as measured using R_1ρ_ and R_2ρ_ NMR experiments). We agree that this is not the only excited state that is possibly present in the system under the conditions used. R_1ρ_ and R_2ρ_ NMR experiments are sensitive to only a range of motions. There are likely other excited states present at pH 6.4 outside the detection window of the timescale sensitive to R_1ρ_ and R_2ρ_ NMR experiments. These additional excited states may also get quenched or enhanced by the lower pH, thereby affecting the monomer population and, hence, its concentration.

The fibrils formed in the two buffer conditions are heterogeneous and have visibly different morphologies. AFM data analysis showed an increased heterogeneity in the fibrils formed at pH 4.6 as compared to that at pH 6.4 (Fig. S2). In addition to the formation of fibrils of similar length (∼0.6 μm) and height (∼12 nm) (fibril state A) at pH 4.6, a significant number of shorted fibrils of length <0.2 μm (fibril state B) were also observed. In addition, the pathway from monomer to fibril may also be different in the two pH conditions, resulting in altered kinetics. It is possible to have perturbations in other excited states at the lower pH, which are not sensed by the relaxation measurements carried out here, resulting in differences in the pathway itself. The data shows that the external perturbation (a decrease in pH) quenches the intrinsic exchange dynamics between M and ES, which leads to an increase in the M population and also enhances fibril formation rates with a new fibril state B. At the lower pH, the ES is quenched, and shaking of the monomer leads to forming an alternate fibril state B (Fig. 6). It remains to be established if the shorter fibrils formed at pH 4.6 are differently toxic to the cells and need to be explored. Thus, the pH stress might work as a switch to turn on/off the exchange dynamics between M and ES to control the fibril formation during physiological and stress conditions. Under normal conditions, ES exists, leading to the formation of fibril state A. However, stress conditions like metabolic perturbation, oxidative stress, etc., alter pH of the cytoplasm, leading to perturbations in the intrinsic conformational exchange between M and ES, thereby affecting the population ratio of M vs ES and increasing the RRM fibril formation in state B.

**Figure 6.**
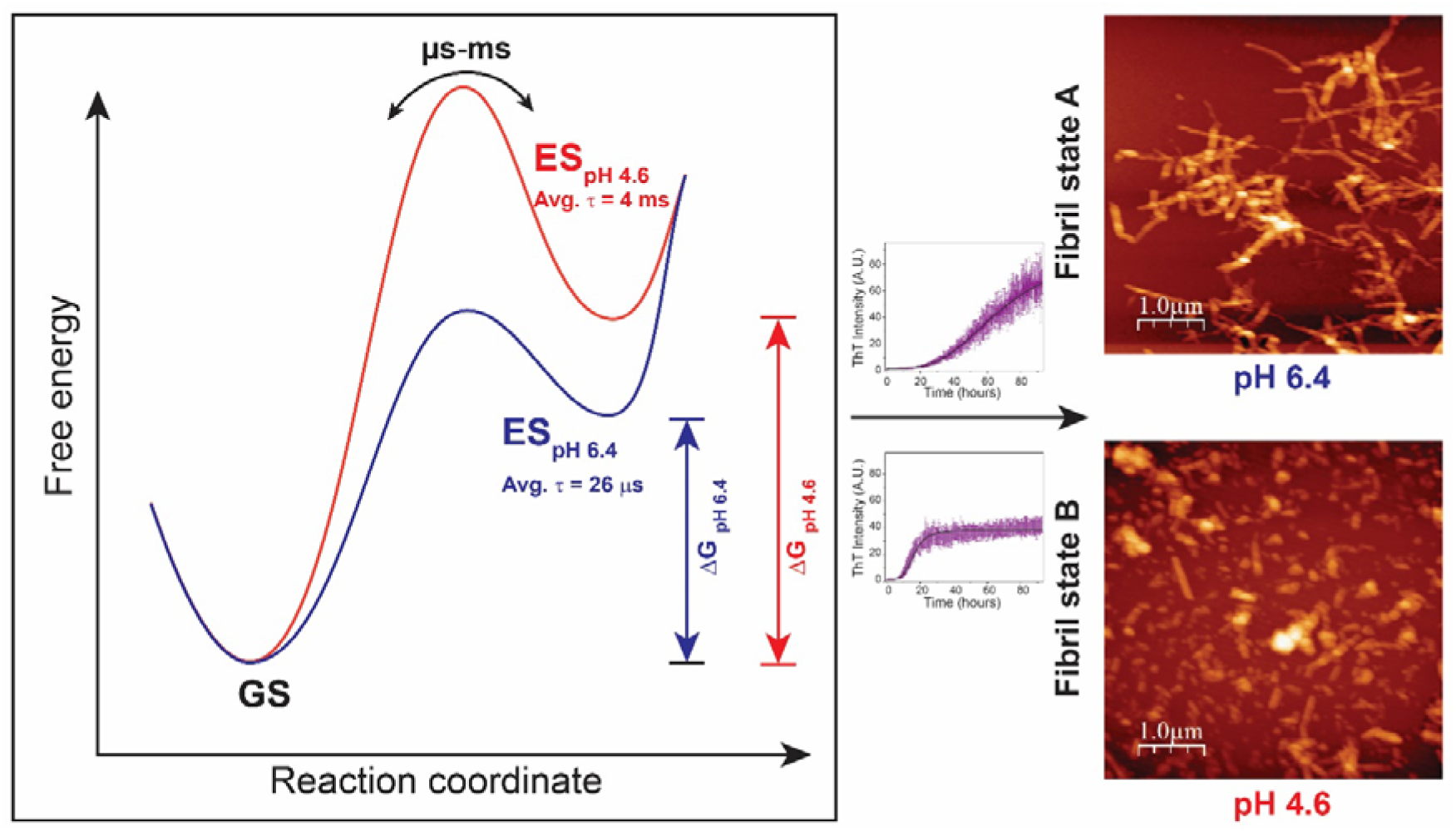
The proposed model for amyloid formation by FUS-RRM and the role of pH stress in destabilizing the ES, causing a change in the fibril state. A cartoon representation of the free energy differences between GS and ES at the two pH values, where corresponding average lifetimes (τ) of ES have been mentioned. A perturbation in the conformational heterogeneity is coupled with the fibril state achieved upon agitation, with differences in the kinetics of fibril formation.

Local pH levels in cells vary differently depending on the type of stress, e.g., temperature perturbation, starvation, metabolic perturbations, osmotic stress, aging and neurodegenerative diseases, cancers, etc [38]. In tumor environments, cellular pH as low as 5.8 has been observed. There is also pH variation across different cellular organelles, ranging from 4.5 to 8.0. While by going down to pH 4.6 *in vitro* conditions in this study, we do not necessarily mimic the exact intracellular low pH values seen in the above-mentioned stresses around FUS condensates, we have tried to show that a low pH can cause a perturbation in the conformational dynamics of a protein thereby perturbing the conformational heterogeneity observed in proteins. Such perturbations to conformational heterogeneity (and not structures and surface charges alone) as a function of pH might also be responsible for the loss or gain of related functions, which in this case is fibril formation by RRM.

Therefore, new strategies can be designed that can shift the population of the excited state(s) towards the higher side, preventing amyloid deposits.

## 5. Conclusion

In this study, we have identified that FUS-RRM undergoes rapid exchange with an excited state (ES), which becomes significantly perturbed at lower pH, leading to a significant change in its average lifetime. Beyond disrupting the conformational heterogeneity of the native state, pH plays a crucial role in accelerating fibril formation kinetics while altering the final fibril state—all without affecting the monomer structure. Given that this conformational heterogeneity appears to be intrinsically linked to both the fibril formation pathway and the resulting fibril state, pH may function as a molecular switch, regulating fibrillation under pathophysiological and stress conditions. Moreover, our findings suggest that structure-based drug discovery efforts, which traditionally focus on native states, should also consider excited-state conformations to expand the target search space for lead compound design. However, this study does not conclude that the identified ES is the only one present. Further investigations using NMR techniques sensitive to a broad range of timescales are required to uncover additional excited states and gain deeper insights into amyloid formation mechanisms. Additionally, other perturbations—such as ATP, RNA, and DNA binding—may also be explored and optimized to perturb the monomer-to-ES exchange process.

## CRediT authorship contribution statement

J.C. and O.A. conceived the study and designed the approach to dynamically characterize the FUS-RRM. O.A. prepared and purified protein samples and measured all the NMR data. O.A. and A.A. analyzed the HARD data. O.A. and L.S. performed all the biophysical experiments. O.A. and J.C. have contributed toward manuscript writing.

## Acknowledgments

The authors acknowledge Prof. Neel Sarovar Bhavesh (ICGEB, New Delhi, India) for the FUS-RRM plasmids. OA acknowledges Prof. Jayant B. Udgaonkar (IISER Pune, India) for providing the facility to measure aggregation kinetics and Mr. Ishtiyaq Ahmad for useful insights into aggregation kinetics. OA also acknowledges Dr. Radha Chauhan (NCCS, India) and Ms. Jyotsana for help with SEC-MALS experiments. OA acknowledges lab members for all the discussion during the writing of this manuscript. The authors acknowledge the High-Field NMR facility at IISER-Pune (co-funded by DST-FIST and IISER Pune). J.C. acknowledges extramural funding from the Science and Engineering Research Board, Govt. of India (EMR/2015/001966 and CRG/2023/002931-C), Department of Biotechnology, Govt. of India (BT/PR24185/BRB/10/1605/2017), Department of Health Research, ICMR (R.11015/14/2023-GIA/HR), and the generous funding from IISER Pune. OA is grateful to DBT-JRF, Govt. of India, for providing a fellowship.

## Data availability

All the raw data has been uploaded to the Mendeley server and is available to download via this link: https://data.mendeley.com/preview/t3cftxm6zb?a=4904313e-e731-4331-8fbd-3ee195aef1eb

