## Supplementary Information for "Investigating the role of conformational heterogeneity in FUS-RRM fibrillation"

Dr. Jeetender Chugh

**Postal address**: C-115, Department of Chemistry, Main Building, Indian Institute of Science Education and Research (IISER), Dr. Homi Bhabha Road, Pashan, Pune 411008, India

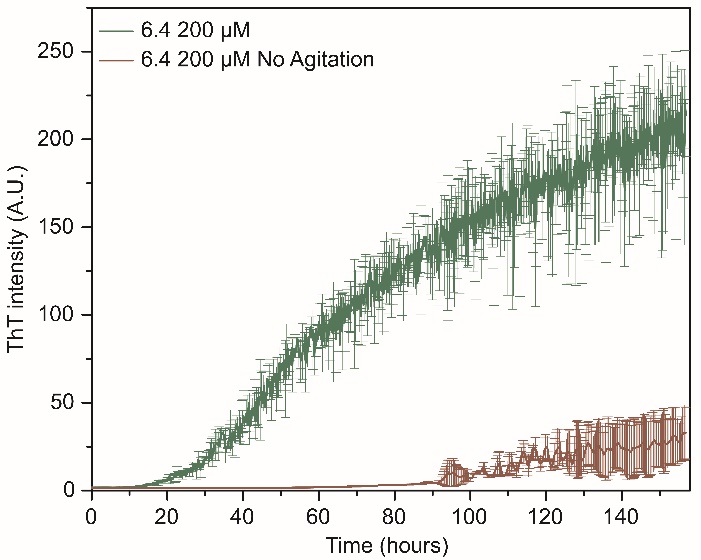

**Figure S1.** Aggregation kinetics at pH 6.4 monitored by ThT intensity while shaking the protein (200 μM) at 60 rpm. Data are represented as mean ± SD of two biological and two technical repeats. ThT intensity at pH 6.4 does not increase significantly when the protein is not agitated. No plateau is achieved even after measurements at a high protein concentration and for 160 hours.

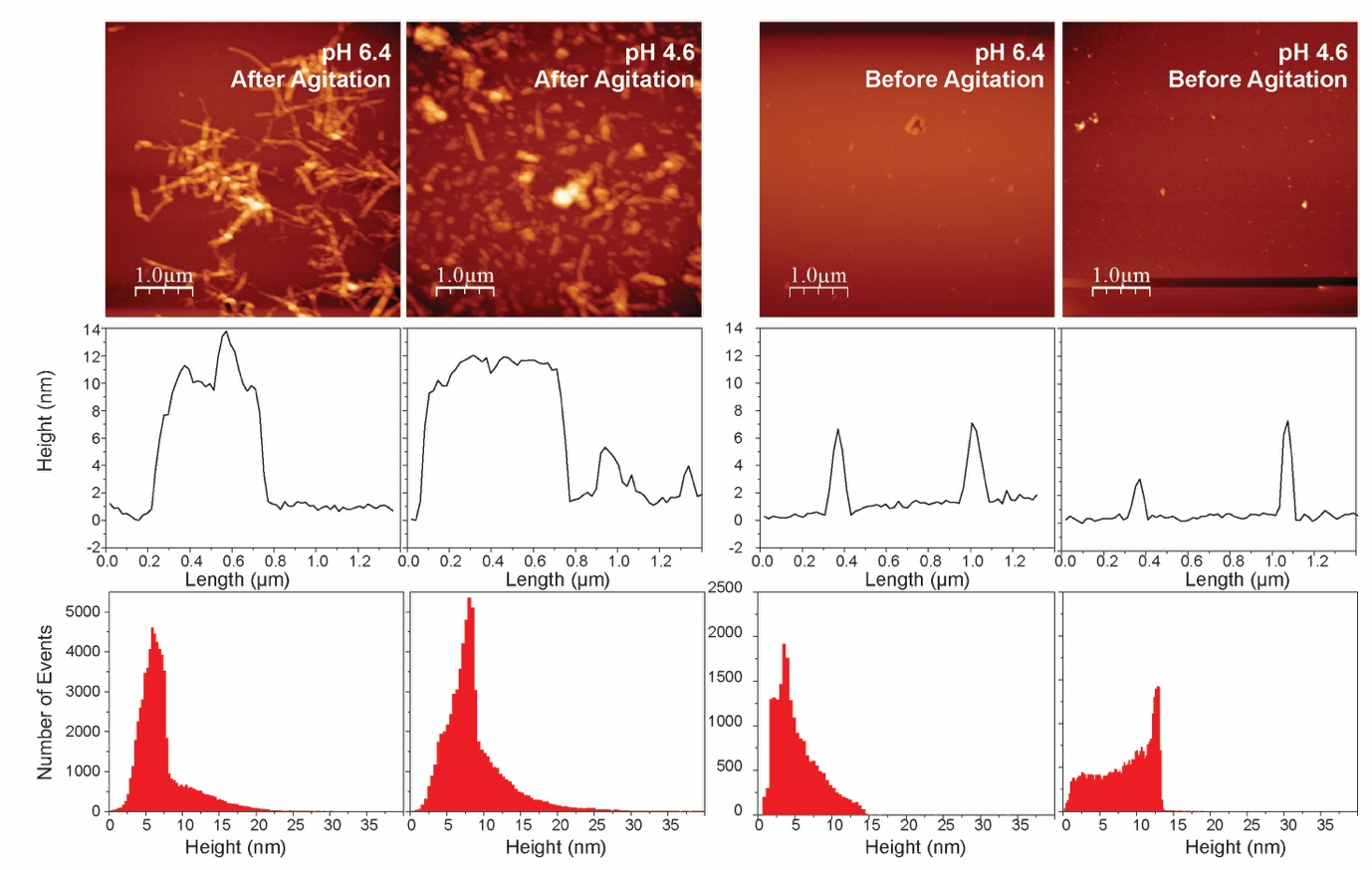

**Figure S2.** AFM measurement on protein samples with and without shaking. Top panel: AFM image of FUS-RRM before and after agitation of the same recorded at pH 6.4 and pH 4.6. Middle panel: Analysis of the corresponding AFM images showing a correlation between height (nm) and length (μm) of the aggregates. Bottom panel: Analysis of the corresponding AFM images showing a correlation between the number of events and height (nm) of the aggregates.

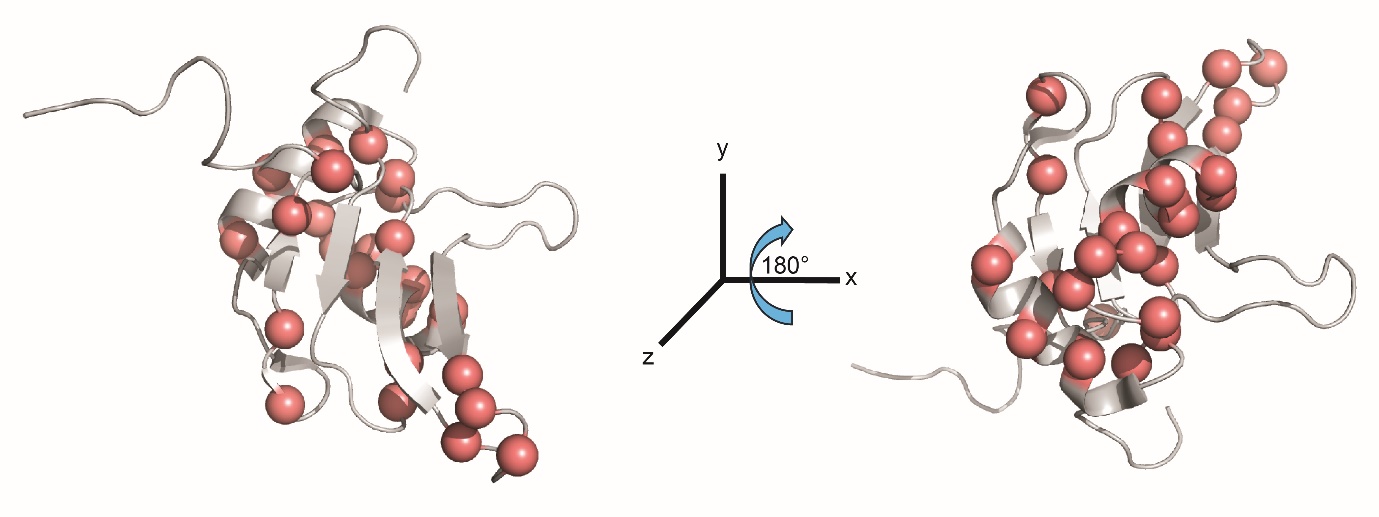

**Figure S3.** The population of the excited state obtained from the HARD NMR experiments mapped on the tertiary structure of the protein. Residues having a population > 5%, shown as spheres, seem to cluster on one face of the protein.

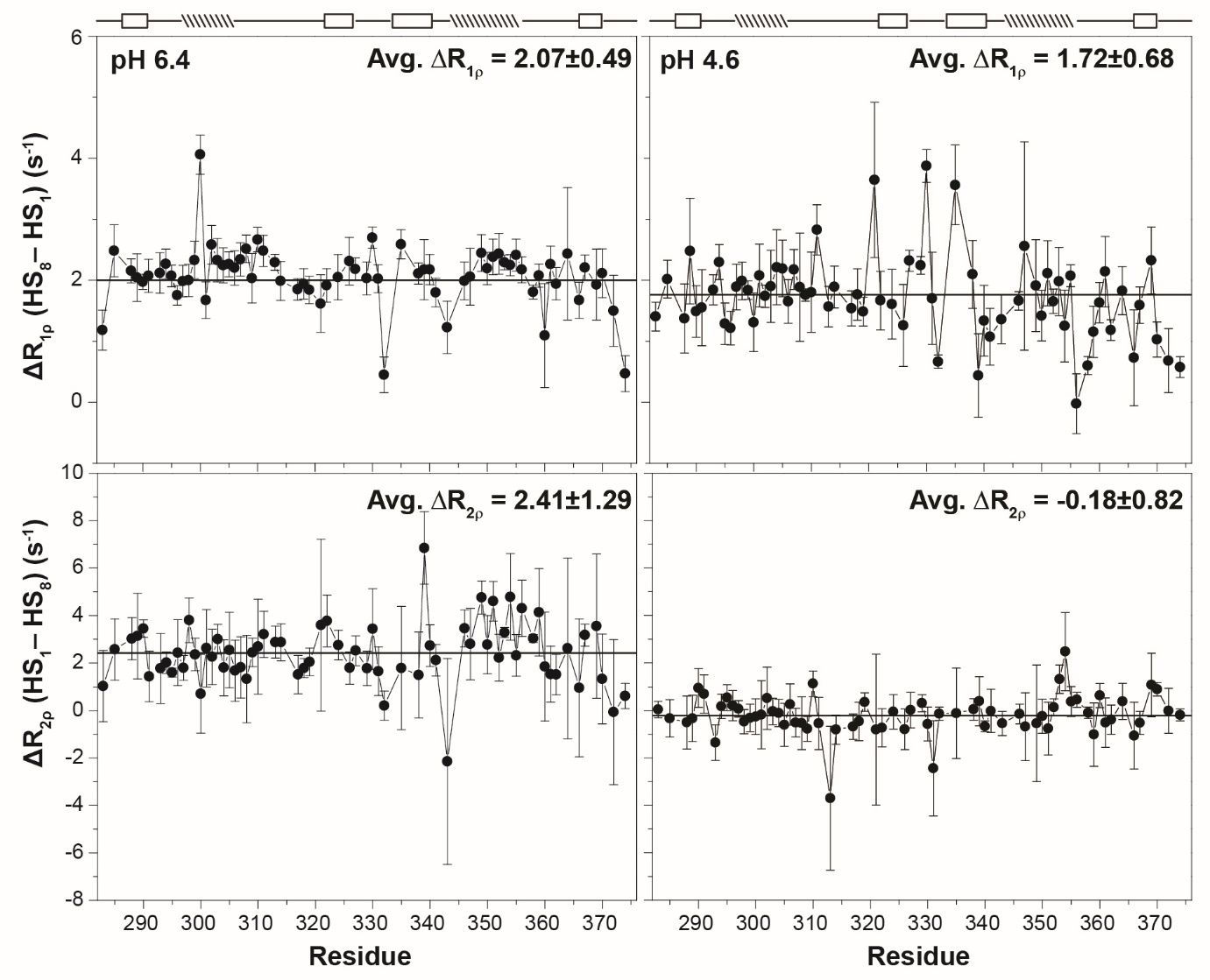

**Figure S4.** ΔR_1ρ_ and ΔR_2ρ_ measured at two stretching factors, HS1 and HS8, of adiabatic pulse to highlight the relaxation dispersion observed at different residues and at two different pH values (6.4 and 4.6). Average values have been marked using a horizontal line. The secondary structure is shown on the top of each panel.

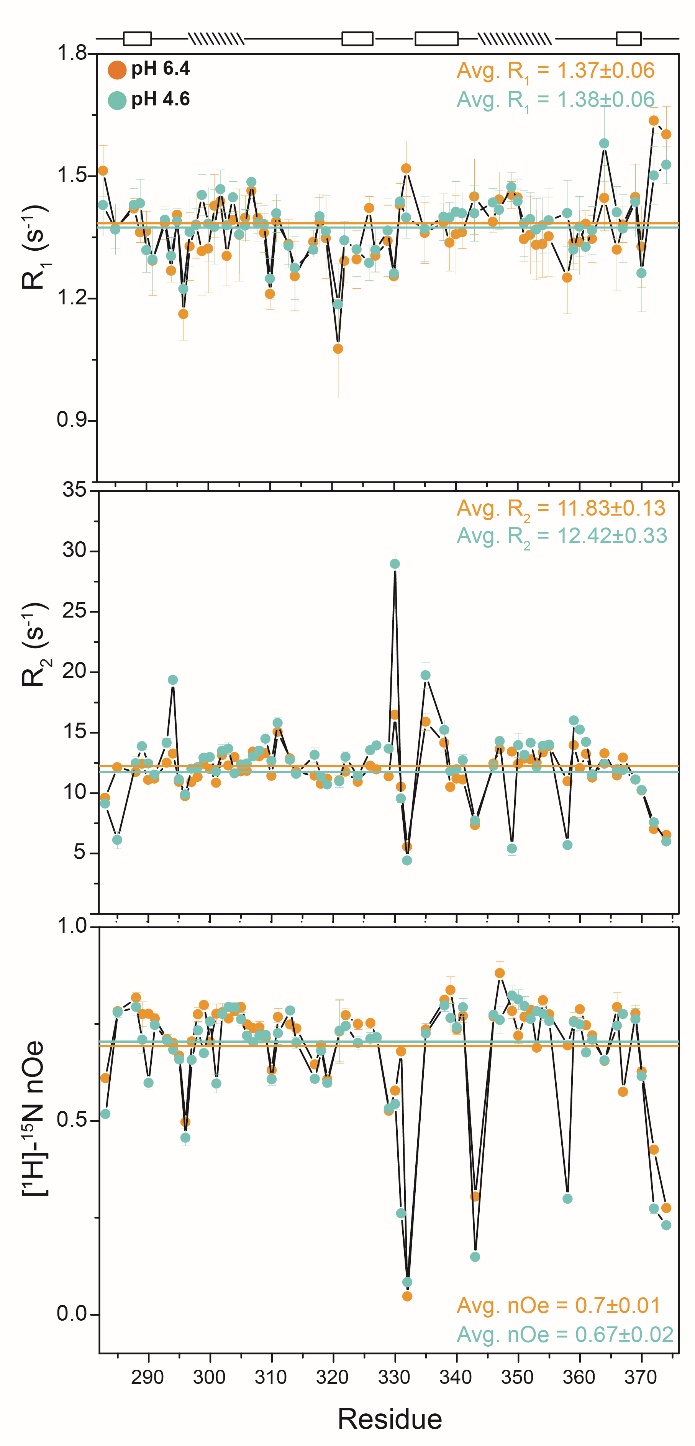

**Figure S5.** Overlay of the R_1_, R_2_, and [^1^H]-^15^N nOe relaxation rates measured for FUS-RRM at pH 6.4 and 4.6. The average line has been shown on the data plots, and average values for the rates have been mentioned in the figure. The secondary structure is shown on the top.

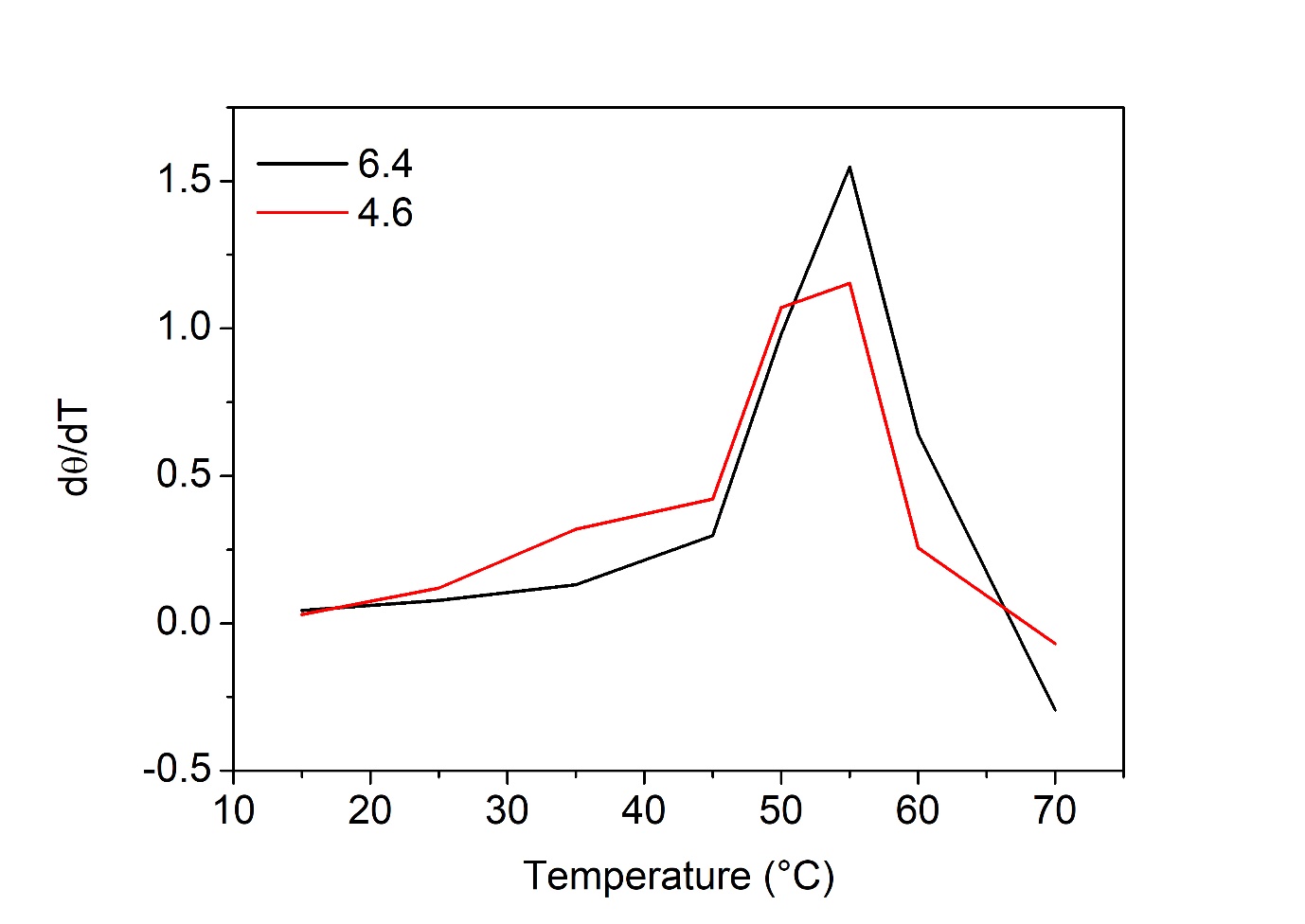

**Figure S6.** T_m_ profile for FUS-RRM for different pH values.

**Table S1**: Lag time (in hours) obtained from fitting the aggregation kinetics data measured at pH 6.4 and 4.6

|  | **pH 6.4** | | | **pH 4.6** | | |
| --- | --- | --- | --- | --- | --- | --- |
|  | 25 µM | 50 µM | 100 µM | 25 µM | 50 µM | 100 µM |
| **Lag time (hours)** | 54.12 | 40.03 | 34.44 | 8.47 | 9.37 | 9.94 |

**Table S2**: Secondary structure estimation from CD spectrum (pH 6.4 and pH 4.6) and the solution structure (pH 7.0).

|  |  | β-strand (%) | α-helix (%) | Loop (%) |
| --- | --- | --- | --- | --- |
| Calculated from PDB ID 2LCW | pH 7.0 | 20.4 | 20.4 | 59.2 |
| Predicted using BeStSel from CD spectrum | pH 6.4 | 19.7 | 14.7 | 65.6 |
|  | pH 4.6 | 21.9 | 11.8 | 66.3 |

**Table S3**: *R_1_* relaxation rates measured from HARD experiment for FUS-RRM (pH 6.4) at 600 MHz NMR spectrometer. Only data for which good fits were obtained have been shown for both the pH values.

| **Residue number** | **R_1_ (s^-1^)** | |
| --- | --- | --- |
|  | **Value** | **Error** |
| D283 | 1.59 | 0.05 |
| N285 | 1.37 | 0.03 |
| F288 | 1.32 | 0.03 |
| V289 | 1.42 | 0.07 |
| Q290 | 1.30 | 0.01 |
| G291 | 1.28 | 0.02 |
| G293 | 1.31 | 0.01 |
| E294 | 1.22 | 0.01 |
| N295 | 1.40 | 0.02 |
| V296 | 1.17 | 0.02 |
| T297 | 1.31 | 0.01 |
| I298 | 1.33 | 0.01 |
| E299 | 1.32 | 0.01 |
| S300 | 1.29 | 0.01 |
| V301 | 1.20 | 0.05 |
| A302 | 1.37 | 0.04 |
| D303 | 1.24 | 0.02 |
| Y304 | 1.37 | 0.02 |
| F305 | 1.28 | 0.01 |
| K306 | 1.42 | 0.03 |
| Q307 | 1.41 | 0.01 |
| I308 | 1.36 | 0.02 |
| G309 | 1.29 | 0.01 |
| I310 | 1.25 | 0.02 |
| I311 | 1.34 | 0.03 |
| T313 | 1.22 | 0.02 |
| N314 | 1.27 | 0.01 |
| T317 | 1.33 | 0.02 |
| G318 | 1.36 | 0.01 |
| Q319 | 1.34 | 0.01 |
| M321 | 1.09 | 0.07 |
| I322 | 1.28 | 0.01 |
| L324 | 1.30 | 0.03 |
| T326 | 1.47 | 0.09 |
| D327 | 1.35 | 0.01 |
| E329 | 1.28 | 0.02 |
| T330 | 1.28 | 0.02 |
| G331 | 1.45 | 0.02 |
| K332 | 1.63 | 0.03 |
| G335 | 1.45 | 0.03 |
| T338 | 1.29 | 0.01 |
| V339 | 1.35 | 0.01 |
| S340 | 1.28 | 0.01 |
| F341 | 1.31 | 0.03 |
| D343 | 1.69 | 0.14 |
| S346 | 1.37 | 0.01 |
| A347 | 1.35 | 0.03 |
| A349 | 1.41 | 0.01 |
| A350 | 1.42 | 0.03 |
| I351 | 1.29 | 0.03 |
| D352 | 1.29 | 0.01 |
| W353 | 1.28 | 0.01 |
| F354 | 1.27 | 0.01 |
| D355 | 1.29 | 0.02 |
| G356 | 1.36 | 0.02 |
| E358 | 1.29 | 0.02 |
| F359 | 1.32 | 0.01 |
| S360 | 1.52 | 0.16 |
| G361 | 1.45 | 0.05 |
| N362 | 1.32 | 0.01 |
| I364 | 1.51 | 0.05 |
| V366 | 1.28 | 0.05 |
| S367 | 1.33 | 0.01 |
| A369 | 1.39 | 0.03 |
| T370 | 1.51 | 0.08 |
| R372 | 1.82 | 0.13 |
| D374 | 1.76 | 0.05 |

**Table S4**: *R_1ρ_* relaxation rates measured using HSn pulses (n=1,2,4,6,8) from HARD experiment for FUS-RRM (pH 6.4) at 600 MHz NMR spectrometer.

| **Residue**  **Number** | ***R_1ρ_* (s^-1^)** | | | | | | | | | |
| --- | --- | --- | --- | --- | --- | --- | --- | --- | --- | --- |
|  | **HS1** | | **HS2** | | **HS4** | | **HS6** | | **HS8** | |
|  | **Value** | **Error** | **Value** | **Error** | **Value** | **Error** | **Value** | **Error** | **Value** | **Error** |
| D283 | 4.21 | 0.28 | 4.41 | 0.05 | 5.03 | 0.10 | 5.39 | 0.28 | 5.39 | 0.16 |
| N285 | 4.22 | 0.18 | 5.21 | 0.19 | 6.10 | 0.29 | 6.47 | 0.42 | 6.70 | 0.39 |
| F288 | 3.85 | 0.19 | 4.74 | 0.13 | 5.60 | 0.17 | 5.92 | 0.21 | 6.00 | 0.07 |
| V289 | 4.16 | 0.21 | 4.51 | 0.07 | 5.39 | 0.33 | 6.13 | 0.20 | 6.20 | 0.33 |
| Q290 | 3.79 | 0.11 | 4.86 | 0.03 | 5.45 | 0.03 | 6.01 | 0.13 | 5.76 | 0.06 |
| G291 | 4.23 | 0.23 | 5.22 | 0.17 | 5.79 | 0.07 | 6.32 | 0.26 | 6.30 | 0.15 |
| G293 | 4.50 | 0.23 | 5.36 | 0.10 | 6.09 | 0.21 | 6.62 | 0.36 | 6.62 | 0.24 |
| E294 | 5.01 | 0.23 | 6.05 | 0.05 | 6.69 | 0.14 | 7.23 | 0.21 | 7.28 | 0.07 |
| N295 | 4.16 | 0.16 | 5.07 | 0.15 | 5.77 | 0.15 | 6.18 | 0.29 | 6.24 | 0.08 |
| V296 | 3.87 | 0.10 | 4.66 | 0.21 | 5.23 | 0.08 | 5.47 | 0.18 | 5.62 | 0.13 |
| T297 | 3.91 | 0.21 | 4.78 | 0.08 | 5.43 | 0.06 | 5.96 | 0.24 | 5.90 | 0.14 |
| I298 | 4.62 | 0.22 | 5.50 | 0.14 | 6.07 | 0.12 | 6.91 | 0.35 | 6.62 | 0.15 |
| E299 | 4.76 | 0.26 | 5.77 | 0.13 | 6.48 | 0.12 | 7.10 | 0.35 | 7.09 | 0.17 |
| S300 | 2.62 | 0.18 | 5.03 | 0.15 | 6.18 | 0.14 | 6.11 | 0.82 | 6.68 | 0.26 |
| V301 | 4.84 | 0.25 | 5.55 | 0.10 | 6.35 | 0.20 | 6.49 | 0.39 | 6.51 | 0.16 |
| A302 | 4.53 | 0.26 | 5.33 | 0.09 | 6.64 | 0.22 | 6.84 | 0.35 | 7.11 | 0.18 |
| D303 | 4.59 | 0.27 | 5.53 | 0.16 | 6.42 | 0.20 | 6.65 | 0.34 | 6.92 | 0.23 |
| Y304 | 4.38 | 0.22 | 5.10 | 0.13 | 5.96 | 0.15 | 6.59 | 0.26 | 6.62 | 0.19 |
| F305 | 4.30 | 0.19 | 5.36 | 0.07 | 6.13 | 0.10 | 6.46 | 0.15 | 6.56 | 0.07 |
| K306 | 4.11 | 0.18 | 4.74 | 0.07 | 5.86 | 0.17 | 5.96 | 0.28 | 6.31 | 0.22 |
| Q307 | 4.90 | 0.23 | 5.89 | 0.15 | 6.71 | 0.10 | 7.19 | 0.29 | 7.24 | 0.14 |
| I308 | 4.20 | 0.20 | 5.28 | 0.18 | 6.49 | 0.25 | 6.68 | 0.33 | 6.72 | 0.10 |
| G309 | 4.60 | 0.35 | 5.38 | 0.17 | 6.05 | 0.11 | 6.74 | 0.43 | 6.63 | 0.21 |
| I310 | 4.25 | 0.12 | 5.30 | 0.32 | 5.88 | 0.11 | 6.34 | 0.20 | 6.91 | 0.17 |
| I311 | 4.58 | 0.17 | 5.79 | 0.25 | 6.50 | 0.20 | 7.18 | 0.26 | 7.06 | 0.19 |
| T313 | 4.09 | 0.11 | 5.04 | 0.12 | 5.97 | 0.19 | 6.17 | 0.14 | 6.38 | 0.08 |
| N314 | 4.49 | 0.28 | 5.33 | 0.14 | 5.98 | 0.10 | 6.49 | 0.32 | 6.48 | 0.15 |
| T317 | 4.22 | 0.15 | 4.94 | 0.09 | 5.51 | 0.07 | 5.97 | 0.28 | 6.07 | 0.09 |
| G318 | 4.25 | 0.23 | 5.12 | 0.13 | 5.81 | 0.09 | 6.34 | 0.23 | 6.19 | 0.10 |
| Q319 | 3.89 | 0.15 | 4.64 | 0.10 | 5.29 | 0.18 | 5.64 | 0.26 | 5.73 | 0.17 |
| M321 | 3.91 | 0.43 | 4.90 | 0.35 | 5.16 | 0.48 | 5.34 | 0.43 | 5.52 | 0.20 |
| I322 | 4.04 | 0.21 | 4.98 | 0.12 | 5.33 | 0.07 | 6.23 | 0.30 | 5.96 | 0.18 |
| L324 | 3.99 | 0.29 | 4.88 | 0.12 | 5.48 | 0.13 | 6.13 | 0.33 | 6.04 | 0.24 |
| T326 | 4.58 | 0.33 | 5.38 | 0.20 | 6.23 | 0.12 | 6.76 | 0.23 | 6.89 | 0.20 |
| D327 | 4.51 | 0.17 | 5.41 | 0.10 | 6.33 | 0.15 | 6.64 | 0.18 | 6.69 | 0.07 |
| E329 | 4.48 | 0.23 | 5.50 | 0.15 | 6.09 | 0.12 | 6.33 | 0.22 | 6.51 | 0.14 |
| T330 | 4.49 | 0.12 | 5.64 | 0.05 | 6.50 | 0.13 | 7.08 | 0.28 | 7.19 | 0.12 |
| G331 | 3.97 | 0.16 | 4.90 | 0.12 | 5.71 | 0.16 | 6.03 | 0.22 | 5.99 | 0.16 |
| K332 | 3.60 | 0.28 | 3.78 | 0.11 | 3.88 | 0.09 | 4.20 | 0.29 | 4.05 | 0.09 |
| G335 | 4.57 | 0.19 | 5.53 | 0.09 | 6.65 | 0.26 | 6.91 | 0.09 | 7.16 | 0.14 |
| T338 | 4.15 | 0.14 | 5.05 | 0.15 | 5.74 | 0.11 | 6.19 | 0.19 | 6.26 | 0.10 |
| V339 | 3.83 | 0.45 | 4.38 | 0.40 | 4.98 | 0.31 | 5.81 | 0.25 | 6.00 | 0.20 |
| S340 | 4.45 | 0.23 | 5.37 | 0.05 | 5.97 | 0.10 | 6.56 | 0.19 | 6.62 | 0.09 |
| F341 | 4.03 | 0.19 | 4.81 | 0.03 | 5.65 | 0.07 | 6.05 | 0.29 | 5.83 | 0.13 |
| D343 | 3.92 | 0.31 | 4.44 | 0.29 | 5.05 | 0.26 | 5.09 | 0.51 | 5.14 | 0.30 |
| S346 | 4.46 | 0.22 | 5.26 | 0.10 | 5.85 | 0.14 | 6.49 | 0.25 | 6.44 | 0.22 |
| A347 | 4.75 | 0.34 | 5.59 | 0.29 | 6.56 | 0.11 | 7.16 | 0.36 | 6.80 | 0.31 |
| A349 | 3.82 | 0.28 | 4.88 | 0.22 | 5.46 | 0.21 | 6.17 | 0.22 | 6.27 | 0.09 |
| A350 | 4.67 | 0.26 | 5.96 | 0.21 | 6.43 | 0.24 | 7.20 | 0.09 | 6.86 | 0.06 |
| I351 | 4.23 | 0.25 | 5.46 | 0.25 | 5.99 | 0.18 | 6.56 | 0.29 | 6.61 | 0.15 |
| D352 | 4.60 | 0.23 | 5.59 | 0.19 | 6.43 | 0.16 | 6.94 | 0.39 | 7.03 | 0.25 |
| W353 | 4.29 | 0.10 | 5.53 | 0.10 | 6.13 | 0.11 | 6.42 | 0.14 | 6.58 | 0.06 |
| F354 | 4.13 | 0.10 | 5.24 | 0.21 | 6.05 | 0.09 | 6.44 | 0.16 | 6.38 | 0.14 |
| D355 | 4.64 | 0.24 | 5.57 | 0.16 | 6.49 | 0.08 | 6.96 | 0.26 | 7.05 | 0.10 |
| G356 | 4.22 | 0.16 | 4.96 | 0.11 | 5.94 | 0.11 | 6.33 | 0.29 | 6.39 | 0.14 |
| E358 | 2.89 | 0.05 | 3.57 | 0.09 | 4.33 | 0.05 | 4.58 | 0.06 | 4.69 | 0.10 |
| F359 | 3.80 | 0.11 | 4.83 | 0.09 | 5.28 | 0.06 | 5.80 | 0.21 | 5.88 | 0.15 |
| S360 | 4.91 | 0.71 | 5.29 | 0.63 | 6.37 | 0.22 | 6.29 | 1.06 | 6.00 | 0.47 |
| G361 | 4.28 | 0.22 | 5.06 | 0.21 | 5.78 | 0.15 | 6.29 | 0.39 | 6.55 | 0.20 |
| N362 | 3.92 | 0.22 | 4.57 | 0.09 | 5.38 | 0.13 | 5.66 | 0.23 | 5.86 | 0.16 |
| I364 | 3.08 | 0.31 | 3.95 | 0.73 | 5.55 | 0.58 | 6.36 | 0.68 | 5.51 | 1.04 |
| V366 | 4.26 | 0.20 | 5.39 | 0.40 | 5.47 | 0.33 | 6.10 | 0.12 | 5.94 | 0.22 |
| S367 | 4.23 | 0.18 | 5.16 | 0.13 | 5.79 | 0.11 | 6.44 | 0.30 | 6.43 | 0.10 |
| A369 | 4.99 | 0.55 | 6.24 | 0.22 | 6.38 | 0.22 | 6.65 | 0.44 | 6.92 | 0.19 |
| T370 | 3.87 | 0.38 | 4.77 | 0.18 | 5.14 | 0.12 | 5.72 | 0.32 | 5.98 | 0.12 |
| R372 | 3.75 | 0.30 | 4.05 | 0.47 | 4.84 | 0.27 | 4.56 | 0.50 | 5.25 | 0.50 |
| D374 | 4.08 | 0.28 | 4.29 | 0.06 | 4.48 | 0.05 | 4.79 | 0.33 | 4.55 | 0.09 |

**Table S5**: *R_2ρ_* relaxation rates measured using HSn pulses (n=1,2,4,6,8) from HARD experiment for FUS-RRM (pH 6.4) at 600 MHz NMR spectrometer.

| **Residue**  **Number** | ***R_2ρ_* (s^-1^)** | | | | | | | | | |
| --- | --- | --- | --- | --- | --- | --- | --- | --- | --- | --- |
|  | **HS1** | | **HS2** | | **HS4** | | **HS6** | | **HS8** | |
|  | **Value** | **Error** | **Value** | **Error** | **Value** | **Error** | **Value** | **Error** | **Value** | **Error** |
| D283 | 11.92 | 0.73 | 11.45 | 0.95 | 12.13 | 1.10 | 10.26 | 1.31 | 10.90 | 1.31 |
| N285 | 14.75 | 1.07 | 15.00 | 0.32 | 14.24 | 0.74 | 11.99 | 0.22 | 12.18 | 0.70 |
| F288 | 13.72 | 0.46 | 13.83 | 0.49 | 13.01 | 1.40 | 10.74 | 1.28 | 10.69 | 0.75 |
| V289 | 15.90 | 1.70 | 14.41 | 1.90 | 15.38 | 0.97 | 13.52 | 1.01 | 12.76 | 0.62 |
| Q290 | 13.99 | 0.26 | 13.32 | 0.33 | 12.63 | 0.23 | 10.98 | 0.54 | 10.53 | 0.25 |
| G291 | 14.10 | 1.05 | 13.41 | 1.35 | 14.10 | 0.65 | 11.88 | 0.42 | 12.66 | 0.14 |
| G293 | 14.52 | 0.84 | 13.93 | 0.66 | 13.73 | 0.78 | 12.32 | 0.95 | 12.74 | 1.21 |
| E294 | 15.73 | 0.43 | 15.88 | 0.32 | 15.82 | 0.18 | 13.42 | 0.43 | 13.71 | 0.15 |
| N295 | 12.53 | 0.17 | 12.36 | 0.38 | 12.44 | 0.64 | 10.37 | 0.50 | 10.93 | 0.07 |
| V296 | 11.71 | 0.94 | 11.41 | 0.59 | 10.46 | 1.94 | 8.25 | 1.35 | 9.27 | 1.03 |
| T297 | 13.11 | 0.50 | 12.62 | 0.79 | 12.79 | 0.40 | 11.13 | 0.40 | 11.31 | 0.15 |
| I298 | 15.53 | 0.54 | 15.21 | 0.46 | 14.05 | 0.88 | 11.67 | 0.93 | 11.73 | 0.77 |
| E299 | 15.15 | 0.41 | 14.87 | 0.35 | 14.55 | 0.52 | 12.43 | 0.43 | 12.78 | 0.54 |
| S300 | 13.49 | 0.82 | 12.41 | 0.71 | 14.18 | 0.68 | 12.31 | 0.50 | 12.79 | 1.45 |
| V301 | 15.44 | 1.33 | 14.85 | 1.63 | 15.15 | 1.36 | 12.80 | 0.60 | 12.82 | 0.94 |
| A302 | 15.08 | 0.91 | 14.75 | 1.12 | 14.34 | 0.55 | 12.43 | 0.50 | 12.82 | 0.72 |
| D303 | 15.29 | 0.41 | 14.99 | 0.26 | 14.56 | 0.87 | 11.98 | 0.19 | 12.29 | 0.47 |
| Y304 | 14.38 | 0.57 | 13.53 | 0.47 | 13.34 | 1.49 | 12.00 | 1.11 | 12.58 | 1.03 |
| F305 | 14.78 | 1.07 | 14.08 | 1.33 | 13.60 | 0.40 | 12.48 | 0.84 | 12.23 | 1.18 |
| K306 | 14.23 | 0.29 | 13.64 | 0.29 | 13.83 | 0.90 | 11.88 | 0.65 | 12.55 | 1.25 |
| Q307 | 14.52 | 0.61 | 14.29 | 0.30 | 13.71 | 0.64 | 12.42 | 0.85 | 12.70 | 1.13 |
| I308 | 13.56 | 1.15 | 12.98 | 0.43 | 12.76 | 1.57 | 11.64 | 1.70 | 12.24 | 1.44 |
| G309 | 14.67 | 0.18 | 14.30 | 0.30 | 14.12 | 0.41 | 12.42 | 0.19 | 12.23 | 0.57 |
| I310 | 14.11 | 1.13 | 13.44 | 0.30 | 13.53 | 0.92 | 11.56 | 1.39 | 11.42 | 1.65 |
| I311 | 16.73 | 0.67 | 15.89 | 0.66 | 16.02 | 0.58 | 13.99 | 0.55 | 13.52 | 0.70 |
| T313 | 14.92 | 0.25 | 14.80 | 0.35 | 14.05 | 0.73 | 12.07 | 0.31 | 12.04 | 0.62 |
| N314 | 14.71 | 0.27 | 14.72 | 0.04 | 14.28 | 0.70 | 11.90 | 0.56 | 11.83 | 0.73 |
| T317 | 13.22 | 0.40 | 13.08 | 0.63 | 12.90 | 1.06 | 11.45 | 0.53 | 11.71 | 0.70 |
| G318 | 13.72 | 0.25 | 13.28 | 0.50 | 13.30 | 0.34 | 11.65 | 0.26 | 11.93 | 0.31 |
| Q319 | 12.42 | 0.36 | 12.06 | 0.25 | 12.05 | 0.55 | 10.35 | 0.34 | 10.37 | 0.46 |
| M321 | 14.60 | 1.33 | 15.99 | 1.18 | 14.73 | 3.13 | 10.44 | 2.63 | 11.00 | 3.37 |
| I322 | 13.92 | 0.97 | 13.17 | 1.09 | 12.06 | 1.25 | 10.44 | 0.28 | 10.15 | 0.52 |
| L324 | 13.32 | 0.52 | 13.13 | 0.42 | 12.51 | 0.32 | 10.48 | 0.68 | 10.57 | 0.35 |
| T326 | 14.95 | 0.52 | 14.26 | 0.75 | 14.57 | 1.21 | 11.82 | 0.64 | 13.15 | 0.44 |
| D327 | 15.13 | 0.14 | 15.02 | 0.28 | 14.51 | 0.90 | 12.48 | 0.54 | 12.61 | 0.61 |
| E329 | 13.13 | 0.40 | 13.08 | 0.14 | 13.01 | 0.32 | 10.88 | 0.20 | 11.35 | 0.61 |
| T330 | 17.69 | 0.81 | 16.78 | 0.33 | 16.37 | 1.00 | 14.26 | 1.26 | 14.26 | 1.50 |
| G331 | 13.56 | 0.48 | 12.79 | 0.64 | 13.49 | 0.96 | 11.54 | 0.36 | 11.91 | 0.91 |
| K332 | 6.46 | 0.32 | 6.56 | 0.24 | 6.90 | 0.44 | 5.53 | 0.26 | 6.26 | 0.53 |
| G335 | 16.63 | 1.68 | 15.84 | 0.80 | 16.68 | 2.19 | 14.58 | 2.07 | 14.84 | 1.97 |
| T338 | 13.95 | 1.17 | 13.80 | 0.74 | 13.85 | 1.37 | 12.62 | 1.02 | 12.46 | 1.38 |
| V339 | 16.34 | 1.39 | 14.77 | 1.81 | 12.26 | 2.00 | 10.73 | 0.48 | 9.49 | 0.63 |
| S340 | 14.36 | 0.05 | 14.19 | 0.21 | 13.18 | 0.91 | 11.43 | 0.60 | 11.63 | 0.87 |
| F341 | 13.51 | 0.52 | 12.80 | 0.82 | 13.45 | 0.99 | 11.75 | 0.50 | 11.40 | 0.41 |
| D343 | 10.83 | 1.50 | 10.68 | 0.71 | 10.78 | 1.73 | 10.88 | 3.03 | 12.97 | 4.08 |
| S346 | 14.99 | 0.39 | 14.39 | 0.25 | 13.90 | 0.34 | 11.82 | 0.64 | 11.53 | 0.67 |
| A347 | 16.69 | 0.79 | 15.69 | 1.28 | 16.17 | 2.33 | 14.42 | 1.61 | 13.89 | 1.28 |
| A349 | 15.07 | 0.64 | 15.65 | 1.01 | 13.77 | 2.16 | 12.22 | 0.83 | 10.31 | 0.28 |
| A350 | 15.75 | 0.95 | 15.76 | 0.83 | 15.79 | 0.34 | 13.02 | 0.81 | 12.97 | 0.77 |
| I351 | 16.76 | 0.80 | 15.73 | 1.25 | 14.41 | 1.83 | 12.14 | 0.52 | 12.15 | 0.22 |
| D352 | 15.28 | 0.66 | 14.92 | 0.41 | 14.74 | 0.66 | 12.56 | 0.59 | 13.05 | 0.73 |
| W353 | 14.72 | 0.17 | 14.69 | 0.12 | 13.87 | 0.45 | 11.72 | 0.39 | 11.44 | 0.13 |
| F354 | 17.20 | 1.11 | 16.63 | 0.48 | 15.97 | 1.08 | 13.08 | 0.98 | 12.41 | 1.45 |
| D355 | 15.98 | 0.39 | 15.77 | 0.20 | 15.15 | 0.71 | 13.10 | 0.56 | 13.66 | 0.79 |
| G356 | 17.11 | 0.61 | 15.98 | 0.29 | 15.72 | 0.86 | 13.00 | 0.72 | 12.80 | 1.02 |
| E358 | 12.68 | 0.09 | 11.77 | 0.08 | 11.34 | 0.56 | 9.76 | 0.33 | 9.64 | 0.14 |
| F359 | 15.50 | 1.16 | 14.29 | 0.53 | 14.01 | 0.98 | 11.29 | 1.31 | 11.37 | 1.44 |
| S360 | 19.34 | 1.23 | 18.11 | 2.06 | 19.07 | 1.37 | 16.45 | 2.28 | 17.49 | 1.94 |
| G361 | 14.40 | 0.84 | 14.31 | 0.18 | 14.71 | 1.31 | 12.62 | 0.65 | 12.88 | 0.83 |
| N362 | 12.30 | 0.35 | 11.73 | 0.06 | 11.82 | 1.03 | 10.48 | 0.67 | 10.78 | 0.77 |
| I364 | 15.01 | 3.47 | 15.22 | 0.93 | 13.53 | 2.33 | 8.17 | 1.07 | 12.39 | 1.59 |
| V366 | 12.85 | 2.06 | 13.05 | 1.42 | 14.14 | 1.45 | 12.11 | 1.79 | 11.90 | 2.05 |
| S367 | 14.48 | 0.44 | 13.82 | 0.44 | 13.67 | 0.69 | 11.56 | 0.42 | 11.30 | 0.14 |
| A369 | 15.03 | 1.01 | 14.31 | 1.24 | 14.19 | 1.87 | 10.72 | 1.29 | 11.47 | 2.86 |
| T370 | 13.29 | 0.83 | 12.62 | 0.78 | 12.83 | 1.71 | 11.26 | 1.60 | 11.96 | 1.71 |
| R372 | 10.53 | 1.46 | 10.41 | 1.81 | 9.87 | 0.58 | 8.76 | 0.74 | 10.60 | 2.70 |
| D374 | 7.73 | 0.32 | 7.63 | 0.14 | 8.00 | 0.34 | 6.26 | 0.31 | 7.12 | 0.43 |

**Table S6**: *R_1_* relaxation rates measured from HARD experiment for FUS-RRM (pH 4.6) at 600 MHz NMR spectrometer.

| **Residue number** | **R_1_ (s^-1^)** | |
| --- | --- | --- |
|  | **Value** | **Error** |
| D283 | 1.53 | 0.03 |
| N285 | 1.52 | 0.07 |
| F288 | 1.77 | 0.03 |
| V289 | 1.50 | 0.11 |
| Q290 | 1.49 | 0.05 |
| G291 | 1.43 | 0.05 |
| G293 | 1.55 | 0.05 |
| E294 | 1.39 | 0.06 |
| N295 | 1.57 | 0.03 |
| V296 | 1.34 | 0.14 |
| T297 | 1.57 | 0.07 |
| I298 | 1.54 | 0.07 |
| E299 | 1.59 | 0.03 |
| S300 | 0.85 | 0.51 |
| V301 | 1.60 | 0.08 |
| A302 | 1.66 | 0.08 |
| D303 | 1.58 | 0.05 |
| Y304 | 1.62 | 0.06 |
| F305 | 1.65 | 0.08 |
| K306 | 1.58 | 0.10 |
| Q307 | 1.65 | 0.05 |
| I308 | 1.72 | 0.11 |
| G309 | 1.53 | 0.06 |
| I310 | 1.42 | 0.07 |
| I311 | 1.66 | 0.09 |
| T313 | 1.55 | 0.04 |
| N314 | 1.52 | 0.05 |
| T317 | 1.49 | 0.05 |
| G318 | 1.56 | 0.08 |
| Q319 | 1.60 | 0.06 |
| M321 | 1.66 | 0.16 |
| I322 | 1.67 | 0.05 |
| L324 | 1.47 | 0.05 |
| T326 | 1.54 | 0.08 |
| D327 | 1.62 | 0.03 |
| E329 | 1.49 | 0.02 |
| T330 | 1.43 | 0.08 |
| G331 | 1.69 | 0.18 |
| K332 | 1.57 | 0.03 |
| G335 | 1.81 | 0.18 |
| T338 | 1.68 | 0.11 |
| V339 | 1.46 | 0.19 |
| S340 | 1.54 | 0.04 |
| F341 | 1.58 | 0.09 |
| D343 | 1.66 | 0.04 |
| S346 | 1.56 | 0.05 |
| A347 | 1.57 | 0.21 |
| A349 | 1.55 | 0.18 |
| A350 | 1.58 | 0.09 |
| I351 | 1.63 | 0.10 |
| D352 | 1.56 | 0.06 |
| W353 | 1.46 | 0.06 |
| F354 | 1.63 | 0.17 |
| D355 | 1.56 | 0.04 |
| G356 | 0.85 | 0.10 |
| E358 | 1.53 | 0.03 |
| F359 | 1.65 | 0.10 |
| S360 | 1.55 | 0.03 |
| G361 | 1.57 | 0.02 |
| N362 | 1.57 | 0.05 |
| I364 | 1.62 | 0.07 |
| V366 | 1.62 | 0.11 |
| S367 | 1.59 | 0.03 |
| A369 | 1.72 | 0.13 |
| T370 | 1.48 | 0.02 |
| R372 | 1.77 | 0.11 |
| D374 | 1.65 | 0.04 |

**Table S7**: *R_1ρ_* relaxation rates measured using HSn pulses (n=1,2,4,6,8) from HARD experiment for FUS-RRM (pH 4.6) at 600 MHz NMR spectrometer.

| **Residue**  **Number** | ***R_1ρ_* (s^-1^)** | | | | | | | | | |
| --- | --- | --- | --- | --- | --- | --- | --- | --- | --- | --- |
|  | **HS1** | | **HS2** | | **HS4** | | **HS6** | | **HS8** | |
|  | **Value** | **Error** | **Value** | **Error** | **Value** | **Error** | **Value** | **Error** | **Value** | **Error** |
| D283 | 3.71 | 0.18 | 4.28 | 0.07 | 4.21 | 0.09 | 4.85 | 0.23 | 5.11 | 0.16 |
| N285 | 4.63 | 0.27 | 5.67 | 0.16 | 5.50 | 0.06 | 6.47 | 0.40 | 6.65 | 0.15 |
| F288 | 4.76 | 0.49 | 4.30 | 0.34 | 5.70 | 0.31 | 5.39 | 0.44 | 6.13 | 0.29 |
| V289 | 4.17 | 0.24 | 4.68 | 0.52 | 6.62 | 0.97 | 6.38 | 0.74 | 6.65 | 0.83 |
| Q290 | 4.54 | 0.19 | 4.75 | 0.22 | 5.16 | 0.31 | 5.95 | 0.17 | 6.03 | 0.38 |
| G291 | 5.06 | 0.34 | 6.35 | 0.33 | 5.51 | 0.17 | 6.87 | 0.48 | 6.61 | 0.53 |
| G293 | 5.10 | 0.21 | 5.76 | 0.35 | 6.11 | 0.19 | 6.75 | 0.34 | 6.94 | 0.13 |
| E294 | 5.56 | 0.19 | 6.42 | 0.07 | 7.11 | 0.12 | 7.94 | 0.24 | 7.86 | 0.21 |
| N295 | 4.55 | 0.30 | 5.46 | 0.13 | 5.10 | 0.07 | 5.82 | 0.20 | 5.84 | 0.17 |
| V296 | 4.87 | 0.23 | 4.19 | 0.77 | 6.29 | 0.38 | 6.26 | 0.41 | 6.09 | 0.15 |
| T297 | 4.14 | 0.08 | 4.73 | 0.21 | 5.21 | 0.15 | 5.72 | 0.35 | 6.03 | 0.37 |
| I298 | 4.67 | 0.21 | 5.49 | 0.43 | 5.53 | 0.29 | 6.79 | 0.36 | 6.66 | 0.22 |
| E299 | 5.15 | 0.08 | 6.04 | 0.13 | 6.09 | 0.11 | 6.82 | 0.16 | 6.99 | 0.13 |
| S300 | 4.72 | 0.13 | 5.16 | 0.52 | 5.94 | 0.46 | 5.81 | 0.38 | 6.03 | 0.46 |
| V301 | 5.34 | 0.28 | 6.19 | 0.19 | 6.44 | 0.42 | 7.51 | 0.67 | 7.42 | 0.43 |
| A302 | 4.62 | 0.11 | 5.44 | 0.26 | 5.90 | 0.52 | 6.94 | 0.59 | 6.36 | 0.14 |
| D303 | 4.67 | 0.19 | 5.27 | 0.16 | 5.53 | 0.21 | 6.43 | 0.41 | 6.57 | 0.56 |
| Y304 | 4.13 | 0.16 | 4.94 | 0.05 | 5.55 | 0.32 | 6.18 | 0.36 | 6.34 | 0.59 |
| F305 | 4.65 | 0.17 | 5.65 | 0.11 | 6.03 | 0.07 | 6.56 | 0.42 | 6.84 | 0.43 |
| K306 | 3.99 | 0.18 | 4.47 | 0.11 | 4.55 | 0.09 | 5.16 | 0.30 | 5.64 | 0.31 |
| Q307 | 5.19 | 0.31 | 6.35 | 0.18 | 6.33 | 0.16 | 7.13 | 0.22 | 7.37 | 0.11 |
| I308 | 4.98 | 0.60 | 6.06 | 0.44 | 5.41 | 0.20 | 5.97 | 0.31 | 6.87 | 0.65 |
| G309 | 4.40 | 0.07 | 5.26 | 0.15 | 5.52 | 0.14 | 6.16 | 0.20 | 6.16 | 0.07 |
| I310 | 5.44 | 0.21 | 6.44 | 0.33 | 5.65 | 0.27 | 6.12 | 0.32 | 7.24 | 0.62 |
| I311 | 4.83 | 0.36 | 6.23 | 0.33 | 7.35 | 0.16 | 7.36 | 0.45 | 7.66 | 0.21 |
| T313 | 4.69 | 0.13 | 4.94 | 0.21 | 5.06 | 0.20 | 5.95 | 0.48 | 6.25 | 0.30 |
| N314 | 4.67 | 0.23 | 5.47 | 0.14 | 5.58 | 0.07 | 6.44 | 0.45 | 6.56 | 0.25 |
| T317 | 5.32 | 0.13 | 5.44 | 0.21 | 5.76 | 0.23 | 6.37 | 0.29 | 6.85 | 0.26 |
| G318 | 4.43 | 0.23 | 4.86 | 0.28 | 5.29 | 0.02 | 5.79 | 0.29 | 6.20 | 0.35 |
| Q319 | 3.93 | 0.17 | 4.77 | 0.20 | 4.71 | 0.09 | 5.20 | 0.23 | 5.42 | 0.17 |
| M321 | 4.65 | 0.71 | 5.59 | 0.35 | 6.81 | 1.12 | 6.98 | 2.17 | 8.29 | 1.05 |
| I322 | 4.31 | 0.42 | 5.25 | 0.31 | 5.29 | 0.32 | 6.22 | 0.34 | 5.97 | 0.24 |
| L324 | 4.33 | 0.34 | 4.77 | 0.35 | 4.46 | 0.25 | 6.10 | 0.38 | 5.94 | 0.46 |
| T326 | 5.79 | 0.56 | 6.51 | 0.37 | 6.46 | 0.44 | 6.92 | 0.28 | 7.05 | 0.37 |
| D327 | 4.62 | 0.16 | 5.16 | 0.19 | 5.77 | 0.15 | 6.82 | 0.21 | 6.94 | 0.06 |
| E329 | 5.51 | 0.06 | 6.61 | 0.27 | 6.54 | 0.17 | 7.09 | 0.09 | 7.76 | 0.13 |
| T330 | 5.52 | 0.15 | 7.26 | 0.10 | 8.81 | 0.29 | 9.87 | 0.08 | 9.39 | 0.22 |
| G331 | 4.81 | 0.22 | 4.33 | 0.97 | 5.24 | 0.35 | 6.10 | 0.35 | 6.51 | 0.72 |
| K332 | 3.37 | 0.10 | 3.60 | 0.06 | 3.37 | 0.02 | 3.80 | 0.23 | 4.04 | 0.05 |
| G335 | 5.59 | 0.22 | 5.74 | 1.05 | 8.09 | 0.50 | 7.77 | 0.66 | 9.15 | 0.62 |
| T338 | 5.01 | 0.50 | 5.97 | 0.40 | 6.59 | 0.26 | 6.62 | 0.29 | 7.11 | 0.24 |
| V339 | 4.76 | 0.29 | 5.19 | 0.51 | 6.60 | 0.41 | 5.92 | 1.26 | 5.20 | 0.62 |
| S340 | 5.58 | 0.40 | 6.32 | 0.40 | 6.15 | 0.45 | 7.38 | 0.25 | 6.92 | 0.41 |
| F341 | 4.63 | 0.25 | 4.70 | 0.21 | 4.44 | 0.20 | 5.68 | 0.43 | 5.70 | 0.39 |
| D343 | 4.35 | 0.37 | 5.17 | 0.42 | 4.91 | 0.31 | 5.22 | 0.25 | 5.70 | 0.16 |
| S346 | 4.65 | 0.13 | 5.09 | 0.14 | 5.29 | 0.15 | 6.35 | 0.32 | 6.31 | 0.07 |
| A347 | 4.48 | 0.75 | 4.68 | 1.03 | 4.83 | 0.16 | 6.27 | 0.90 | 7.05 | 1.53 |
| A349 | 4.34 | 0.48 | 5.55 | 0.24 | 6.01 | 1.11 | 7.72 | 0.41 | 6.25 | 0.58 |
| A350 | 5.67 | 0.18 | 6.38 | 0.36 | 7.05 | 0.17 | 8.25 | 0.75 | 7.09 | 0.38 |
| I351 | 4.52 | 0.06 | 5.63 | 0.32 | 6.28 | 0.47 | 6.41 | 0.46 | 6.63 | 0.53 |
| D352 | 5.27 | 0.13 | 6.14 | 0.26 | 6.03 | 0.07 | 6.50 | 0.16 | 6.92 | 0.16 |
| W353 | 5.32 | 0.46 | 6.56 | 0.33 | 6.75 | 0.39 | 7.00 | 0.19 | 7.30 | 0.31 |
| F354 | 4.86 | 0.30 | 5.66 | 0.29 | 5.89 | 0.25 | 6.73 | 0.63 | 6.11 | 0.50 |
| D355 | 4.99 | 0.10 | 5.69 | 0.11 | 6.12 | 0.17 | 7.00 | 0.18 | 7.07 | 0.15 |
| G356 | 2.85 | 0.37 | 3.12 | 0.22 | 3.12 | 0.06 | 3.39 | 0.46 | 2.83 | 0.32 |
| E358 | 2.86 | 0.10 | 3.06 | 0.14 | 2.90 | 0.14 | 3.26 | 0.21 | 3.46 | 0.11 |
| F359 | 4.28 | 0.18 | 4.85 | 0.20 | 4.81 | 0.36 | 5.43 | 0.25 | 5.43 | 0.38 |
| S360 | 5.71 | 0.32 | 6.40 | 0.45 | 6.64 | 0.29 | 7.60 | 0.29 | 7.34 | 0.08 |
| G361 | 4.40 | 0.24 | 5.02 | 0.41 | 5.06 | 0.17 | 5.70 | 0.21 | 6.54 | 0.52 |
| N362 | 3.96 | 0.11 | 4.79 | 0.04 | 4.62 | 0.05 | 5.18 | 0.23 | 5.14 | 0.13 |
| I364 | 5.06 | 0.19 | 5.62 | 0.40 | 5.97 | 0.54 | 6.92 | 0.34 | 6.89 | 0.34 |
| V366 | 5.38 | 0.62 | 5.86 | 0.55 | 7.12 | 0.63 | 7.15 | 0.28 | 6.11 | 0.49 |
| S367 | 4.56 | 0.23 | 5.21 | 0.16 | 5.38 | 0.17 | 5.97 | 0.48 | 6.15 | 0.19 |
| A369 | 4.33 | 0.48 | 5.63 | 0.44 | 6.29 | 0.57 | 7.25 | 0.71 | 6.66 | 0.27 |
| T370 | 4.25 | 0.18 | 4.49 | 0.23 | 4.55 | 0.08 | 5.20 | 0.34 | 5.28 | 0.22 |
| R372 | 3.65 | 0.27 | 3.93 | 0.16 | 3.78 | 0.42 | 4.44 | 0.42 | 4.33 | 0.45 |
| D374 | 4.59 | 0.16 | 4.82 | 0.23 | 4.60 | 0.17 | 4.87 | 0.05 | 5.17 | 0.05 |

**Table S8**: *R_2ρ_* relaxation rates measured using HSn pulses (n=1,2,4,6,8) from HARD experiment for FUS-RRM (pH 4.6) at 600 MHz NMR spectrometer.

| **Residue**  **Number** | ***R_2ρ_* (s^-1^)** | | | | | | | | | |
| --- | --- | --- | --- | --- | --- | --- | --- | --- | --- | --- |
|  | **HS1** | | **HS2** | | **HS4** | | **HS6** | | **HS8** | |
|  | **Value** | **Error** | **Value** | **Error** | **Value** | **Error** | **Value** | **Error** | **Value** | **Error** |
| D283 | 9.47 | 0.16 | 9.61 | 0.20 | 10.13 | 0.13 | 9.07 | 0.18 | 9.43 | 0.30 |
| N285 | 13.36 | 0.31 | 13.98 | 0.42 | 13.49 | 0.17 | 12.62 | 0.33 | 13.70 | 0.73 |
| F288 | 13.30 | 0.36 | 11.99 | 0.92 | 13.70 | 0.79 | 12.45 | 0.43 | 13.81 | 1.06 |
| V289 | 13.47 | 0.79 | 13.80 | 0.64 | 14.38 | 0.44 | 12.55 | 1.22 | 13.81 | 0.58 |
| Q290 | 13.60 | 0.51 | 13.10 | 0.39 | 13.21 | 0.45 | 11.28 | 0.74 | 12.66 | 0.64 |
| G291 | 13.11 | 0.40 | 13.22 | 0.84 | 13.41 | 0.13 | 12.48 | 0.49 | 12.41 | 0.70 |
| G293 | 14.20 | 0.24 | 13.89 | 0.78 | 14.47 | 0.32 | 13.25 | 0.31 | 15.56 | 0.70 |
| E294 | 19.17 | 0.32 | 19.16 | 0.60 | 18.70 | 0.30 | 17.90 | 0.55 | 19.00 | 0.41 |
| N295 | 11.63 | 0.21 | 11.81 | 0.50 | 11.54 | 0.60 | 11.33 | 0.19 | 11.09 | 0.51 |
| V296 | 9.45 | 0.62 | 10.00 | 0.63 | 8.89 | 0.61 | 10.57 | 0.31 | 9.25 | 0.14 |
| T297 | 12.47 | 0.07 | 13.26 | 0.39 | 12.99 | 0.51 | 12.60 | 0.28 | 12.39 | 0.33 |
| I298 | 12.85 | 0.41 | 13.93 | 0.34 | 14.00 | 0.22 | 12.29 | 0.65 | 13.32 | 0.30 |
| E299 | 13.48 | 0.12 | 14.24 | 0.25 | 13.85 | 0.32 | 13.56 | 0.31 | 13.79 | 0.57 |
| S300 | 12.93 | 0.21 | 13.89 | 0.22 | 13.12 | 0.22 | 13.82 | 1.15 | 13.19 | 0.72 |
| V301 | 11.22 | 1.28 | 12.20 | 1.00 | 12.20 | 1.17 | 12.55 | 0.63 | 11.40 | 0.66 |
| A302 | 14.27 | 0.48 | 12.99 | 0.49 | 14.06 | 0.25 | 12.67 | 0.75 | 13.76 | 1.23 |
| D303 | 13.59 | 0.26 | 13.48 | 0.49 | 13.35 | 0.38 | 12.63 | 0.39 | 13.63 | 0.50 |
| Y304 | 12.92 | 0.28 | 12.88 | 0.30 | 13.40 | 0.54 | 12.06 | 0.38 | 13.03 | 0.50 |
| F305 | 12.29 | 0.43 | 13.66 | 0.82 | 13.78 | 0.32 | 12.62 | 0.37 | 12.90 | 0.80 |
| K306 | 13.10 | 0.38 | 12.56 | 0.38 | 13.30 | 0.44 | 12.34 | 0.39 | 12.85 | 0.79 |
| Q307 | 13.21 | 0.03 | 14.00 | 0.40 | 13.48 | 0.38 | 13.08 | 0.19 | 13.73 | 0.61 |
| I308 | 13.32 | 0.34 | 13.22 | 0.36 | 13.24 | 0.22 | 12.55 | 0.48 | 13.86 | 1.06 |
| G309 | 12.74 | 0.32 | 13.16 | 0.47 | 13.14 | 0.25 | 13.00 | 0.36 | 13.52 | 0.42 |
| I310 | 12.35 | 0.23 | 12.78 | 0.26 | 11.58 | 0.54 | 12.01 | 0.30 | 11.22 | 0.48 |
| I311 | 16.55 | 0.62 | 17.17 | 1.03 | 17.33 | 0.78 | 16.42 | 1.12 | 17.10 | 0.90 |
| T313 | 9.32 | 2.98 | 13.58 | 0.54 | 12.60 | 0.54 | 12.39 | 0.57 | 13.02 | 0.53 |
| N314 | 13.07 | 0.24 | 13.10 | 0.32 | 13.84 | 0.24 | 12.81 | 0.21 | 13.88 | 0.57 |
| T317 | 12.14 | 0.42 | 12.86 | 0.62 | 12.83 | 0.23 | 12.04 | 0.42 | 12.83 | 0.33 |
| G318 | 12.07 | 0.34 | 12.59 | 0.19 | 12.90 | 0.40 | 12.02 | 0.39 | 12.54 | 0.74 |
| Q319 | 10.97 | 0.12 | 10.83 | 0.53 | 11.01 | 0.23 | 10.19 | 0.11 | 10.62 | 0.37 |
| M321 | 14.69 | 2.55 | 12.78 | 1.90 | 16.52 | 1.58 | 19.67 | 1.95 | 15.50 | 1.89 |
| I322 | 11.96 | 0.74 | 12.70 | 0.55 | 12.88 | 0.30 | 11.35 | 0.56 | 12.69 | 0.33 |
| L324 | 11.87 | 0.47 | 11.58 | 0.36 | 12.20 | 0.15 | 10.92 | 0.33 | 11.92 | 0.55 |
| T326 | 13.03 | 0.42 | 15.39 | 0.57 | 14.63 | 0.34 | 13.23 | 0.15 | 13.82 | 0.75 |
| D327 | 15.76 | 0.58 | 15.15 | 0.74 | 16.22 | 0.37 | 14.17 | 0.30 | 15.75 | 0.48 |
| E329 | 13.76 | 0.11 | 14.77 | 0.57 | 13.99 | 0.36 | 12.90 | 0.36 | 13.45 | 0.34 |
| T330 | 25.32 | 0.51 | 24.90 | 0.66 | 24.57 | 0.29 | 25.92 | 0.32 | 25.89 | 0.50 |
| G331 | 11.87 | 0.85 | 13.41 | 1.16 | 12.91 | 0.78 | 14.90 | 1.13 | 14.31 | 1.82 |
| K332 | 4.68 | 0.06 | 4.99 | 0.20 | 5.54 | 0.13 | 4.77 | 0.17 | 4.82 | 0.27 |
| G335 | 19.85 | 1.55 | 20.22 | 1.26 | 19.83 | 2.20 | 18.38 | 2.43 | 19.97 | 1.11 |
| T338 | 13.98 | 0.22 | 15.47 | 0.80 | 13.39 | 0.89 | 12.72 | 1.22 | 13.93 | 0.40 |
| V339 | 12.26 | 0.51 | 12.93 | 0.35 | 13.18 | 0.52 | 12.24 | 0.64 | 11.88 | 0.90 |
| S340 | 13.23 | 0.22 | 14.25 | 0.78 | 13.89 | 0.70 | 13.04 | 0.29 | 13.89 | 0.06 |
| F341 | 12.33 | 0.77 | 12.73 | 0.40 | 13.86 | 0.42 | 11.83 | 0.56 | 12.35 | 0.49 |
| D343 | 6.86 | 0.03 | 7.86 | 0.25 | 8.05 | 0.20 | 7.11 | 0.12 | 7.41 | 0.50 |
| S346 | 12.40 | 0.16 | 12.71 | 0.16 | 12.43 | 0.22 | 12.16 | 0.24 | 12.55 | 0.39 |
| A347 | 16.04 | 1.21 | 16.92 | 2.10 | 18.69 | 0.41 | 15.42 | 1.12 | 16.73 | 0.76 |
| A349 | 12.53 | 0.60 | 14.24 | 1.21 | 13.89 | 0.80 | 13.20 | 0.97 | 13.08 | 2.38 |
| A350 | 13.47 | 0.67 | 13.97 | 0.46 | 13.68 | 0.46 | 12.98 | 0.72 | 13.72 | 0.26 |
| I351 | 14.24 | 0.63 | 15.29 | 0.31 | 15.78 | 0.64 | 13.36 | 0.60 | 15.01 | 0.92 |
| D352 | 15.02 | 0.32 | 15.57 | 0.44 | 14.91 | 0.50 | 14.37 | 0.46 | 14.89 | 0.17 |
| W353 | 13.36 | 0.42 | 13.91 | 0.76 | 14.64 | 0.80 | 12.89 | 0.32 | 12.05 | 0.43 |
| F354 | 18.53 | 1.14 | 18.26 | 1.02 | 17.79 | 0.37 | 19.92 | 2.08 | 16.05 | 1.19 |
| D355 | 14.48 | 0.18 | 14.08 | 0.13 | 13.92 | 0.15 | 14.11 | 0.20 | 14.10 | 0.61 |
| G356 | 6.51 | 0.25 | 6.42 | 0.43 | 7.33 | 0.51 | 5.97 | 0.19 | 6.06 | 0.20 |
| E358 | 4.96 | 0.11 | 5.09 | 0.25 | 5.72 | 0.20 | 5.12 | 0.12 | 5.06 | 0.21 |
| F359 | 14.83 | 0.39 | 15.59 | 0.21 | 14.86 | 0.19 | 15.80 | 0.35 | 15.85 | 1.28 |
| S360 | 16.88 | 0.33 | 17.81 | 0.53 | 16.12 | 0.83 | 16.66 | 0.22 | 16.25 | 0.38 |
| G361 | 15.38 | 0.27 | 14.99 | 0.40 | 15.73 | 0.52 | 15.87 | 0.74 | 15.89 | 1.01 |
| N362 | 10.66 | 0.17 | 10.64 | 0.35 | 11.30 | 0.34 | 10.38 | 0.25 | 11.05 | 0.59 |
| I364 | 12.66 | 0.70 | 14.39 | 0.82 | 13.25 | 0.88 | 13.20 | 0.49 | 12.28 | 0.31 |
| V366 | 13.67 | 0.94 | 14.17 | 0.84 | 12.13 | 0.90 | 13.38 | 1.75 | 14.73 | 1.05 |
| S367 | 12.33 | 0.30 | 12.79 | 0.30 | 12.76 | 0.31 | 11.93 | 0.27 | 12.86 | 0.38 |
| A369 | 13.80 | 1.21 | 13.19 | 0.65 | 12.95 | 0.57 | 13.04 | 0.42 | 12.73 | 0.58 |
| T370 | 11.55 | 0.22 | 11.38 | 0.57 | 11.57 | 0.24 | 10.65 | 0.40 | 10.66 | 0.18 |
| R372 | 6.36 | 0.21 | 7.22 | 0.50 | 7.91 | 0.42 | 7.34 | 0.49 | 6.37 | 0.93 |
| D374 | 5.64 | 0.08 | 6.61 | 0.35 | 6.47 | 0.20 | 5.82 | 0.11 | 5.84 | 0.24 |

**Table S9**: R_1_, R_2_, [^1^H]-^15^N nOe relaxation rates measured for FUS-RRM (pH 6.4) at 600 MHz NMR spectrometer.

| **Residue number** | **R_1_ (s^-1^)** | | **R_2_ (s^-1^)** | | **[^1^H]-^15^N nOe** | |
| --- | --- | --- | --- | --- | --- | --- |
|  | **Value** | **Error** | **Value** | **Error** | **Value** | **Error** |
| D283 | 1.51 | 0.06 | 9.58 | 0.06 | 0.61 | 0.01 |
| N285 | 1.37 | 0.05 | 12.13 | 0.06 | 0.78 | 0.01 |
| F288 | 1.42 | 0.03 | 11.73 | 0.07 | 0.82 | 0.01 |
| V289 | 1.36 | 0.02 | 12.41 | 0.31 | 0.78 | 0.03 |
| Q290 | 1.36 | 0.05 | 11.08 | 0.09 | 0.78 | 0.01 |
| G291 | 1.29 | 0.08 | 11.18 | 0.16 | 0.76 | 0.01 |
| G293 | 1.39 | 0.04 | 12.49 | 0.05 | 0.71 | 0.01 |
| E294 | 1.27 | 0.03 | 13.24 | 0.05 | 0.70 | 0.01 |
| N295 | 1.40 | 0.01 | 10.91 | 0.04 | 0.67 | 0.01 |
| V296 | 1.16 | 0.06 | 9.73 | 0.07 | 0.50 | 0.02 |
| T297 | 1.33 | 0.08 | 10.90 | 0.07 | 0.70 | 0.01 |
| I298 | 1.38 | 0.08 | 11.33 | 0.10 | 0.77 | 0.02 |
| E299 | 1.32 | 0.11 | 12.40 | 0.11 | 0.80 | 0.01 |
| S300 | 1.32 | 0.11 | 11.97 | 0.11 | 0.70 | 0.01 |
| V301 | 1.43 | 0.08 | 10.83 | 0.13 | 0.78 | 0.03 |
| A302 | 1.44 | 0.03 | 13.13 | 0.23 | 0.78 | 0.02 |
| D303 | 1.30 | 0.07 | 12.27 | 0.04 | 0.76 | 0.01 |
| Y304 | 1.39 | 0.02 | 12.97 | 0.07 | 0.78 | 0.01 |
| F305 | 1.36 | 0.11 | 11.77 | 0.09 | 0.79 | 0.01 |
| K306 | 1.40 | 0.05 | 11.83 | 0.07 | 0.75 | 0.01 |
| Q307 | 1.46 | 0.01 | 13.42 | 0.09 | 0.74 | 0.01 |
| I308 | 1.40 | 0.04 | 13.03 | 0.13 | 0.74 | 0.02 |
| G309 | 1.36 | 0.05 | 13.27 | 0.08 | 0.71 | 0.01 |
| I310 | 1.21 | 0.04 | 11.40 | 0.09 | 0.63 | 0.01 |
| I311 | 1.39 | 0.05 | 15.05 | 0.18 | 0.77 | 0.02 |
| T313 | 1.33 | 0.06 | 12.86 | 0.11 | 0.75 | 0.01 |
| N314 | 1.25 | 0.08 | 11.76 | 0.09 | 0.74 | 0.01 |
| T317 | 1.34 | 0.03 | 11.43 | 0.11 | 0.65 | 0.01 |
| G318 | 1.39 | 0.06 | 10.76 | 0.10 | 0.70 | 0.01 |
| Q319 | 1.35 | 0.10 | 11.19 | 0.04 | 0.61 | 0.01 |
| M321 | 1.08 | 0.12 | 11.03 | 0.28 | 0.73 | 0.08 |
| I322 | 1.29 | 0.05 | 11.76 | 0.15 | 0.77 | 0.02 |
| L324 | 1.30 | 0.07 | 10.92 | 0.11 | 0.75 | 0.01 |
| T326 | 1.42 | 0.03 | 12.26 | 0.17 | 0.75 | 0.01 |
| D327 | 1.30 | 0.04 | 11.94 | 0.09 | 0.71 | 0.01 |
| E329 | 1.34 | 0.09 | 11.37 | 0.04 | 0.53 | 0.004 |
| T330 | 1.25 | 0.02 | 16.46 | 0.17 | 0.58 | 0.01 |
| G331 | 1.43 | 0.05 | 10.51 | 0.09 | 0.68 | 0.01 |
| K332 | 1.52 | 0.07 | 5.53 | 0.04 | 0.05 | 0.01 |
| G335 | 1.36 | 0.07 | 15.90 | 0.71 | 0.74 | 0.02 |
| T338 | 1.39 | 0.05 | 14.15 | 0.76 | 0.81 | 0.01 |
| V339 | 1.34 | 0.07 | 10.48 | 0.13 | 0.84 | 0.03 |
| S340 | 1.36 | 0.10 | 11.19 | 0.09 | 0.74 | 0.01 |
| F341 | 1.36 | 0.04 | 11.10 | 0.11 | 0.77 | 0.02 |
| D343 | 1.45 | 0.09 | 7.34 | 0.09 | 0.30 | 0.01 |
| S346 | 1.39 | 0.02 | 12.43 | 0.09 | 0.77 | 0.01 |
| A347 | 1.44 | 0.07 | 13.64 | 0.19 | 0.88 | 0.03 |
| A349 | 1.45 | 0.03 | 13.42 | 0.13 | 0.78 | 0.02 |
| A350 | 1.45 | 0.02 | 12.39 | 0.19 | 0.72 | 0.02 |
| I351 | 1.35 | 0.05 | 12.84 | 0.15 | 0.77 | 0.02 |
| D352 | 1.36 | 0.10 | 12.78 | 0.13 | 0.79 | 0.01 |
| W353 | 1.33 | 0.09 | 12.51 | 0.07 | 0.69 | 0.01 |
| F354 | 1.33 | 0.08 | 13.36 | 0.11 | 0.81 | 0.01 |
| D355 | 1.35 | 0.10 | 13.88 | 0.03 | 0.77 | 0.01 |
| E358 | 1.25 | 0.09 | 10.96 | 0.05 | 0.69 | 0.01 |
| F359 | 1.34 | 0.05 | 13.94 | 0.14 | 0.76 | 0.02 |
| S360 | 1.34 | 0.08 | 12.03 | 0.25 | 0.79 | 0.02 |
| G361 | 1.38 | 0.03 | 13.24 | 0.12 | 0.75 | 0.01 |
| N362 | 1.35 | 0.06 | 11.26 | 0.03 | 0.72 | 0.01 |
| I364 | 1.45 | 0.08 | 13.28 | 0.06 | 0.65 | 0.01 |
| V366 | 1.32 | 0.10 | 11.46 | 0.26 | 0.79 | 0.04 |
| S367 | 1.38 | 0.04 | 12.92 | 0.08 | 0.57 | 0.01 |
| A369 | 1.45 | 0.08 | 11.10 | 0.12 | 0.78 | 0.02 |
| T370 | 1.33 | 0.10 | 10.19 | 0.10 | 0.63 | 0.01 |
| R372 | 1.64 | 0.03 | 7.00 | 0.09 | 0.43 | 0.02 |
| D374 | 1.60 | 0.07 | 6.51 | 0.06 | 0.27 | 0.003 |

**Table S10**: R_1_, R_2_, [^1^H]-^15^N nOe relaxation rates measured for FUS-RRM (pH 4.6) at 600 MHz NMR spectrometer.

| **Residue number** | **R_1_ (s^-1^)** | | **R_2_ (s^-1^)** | | **[^1^H]-^15^N nOe** | |
| --- | --- | --- | --- | --- | --- | --- |
|  | **Value** | **Error** | **Value** | **Error** | **Value** | **Error** |
| D283 | 1.43 | 0.08 | 9.12 | 0.07 | 0.52 | 0.01 |
| N285 | 1.37 | 0.06 | 6.11 | 0.71 | 0.78 | 0.01 |
| F288 | 1.43 | 0.04 | 12.44 | 0.57 | 0.79 | 0.02 |
| V289 | 1.43 | 0.06 | 13.86 | 0.27 | 0.71 | 0.04 |
| Q290 | 1.32 | 0.05 | 12.42 | 0.27 | 0.60 | 0.02 |
| G291 | 1.29 | 0.08 | 11.51 | 0.32 | 0.75 | 0.02 |
| G293 | 1.39 | 0.03 | 14.14 | 0.38 | 0.71 | 0.01 |
| E294 | 1.30 | 0.05 | 19.35 | 0.21 | 0.68 | 0.01 |
| N295 | 1.39 | 0.03 | 11.13 | 0.16 | 0.66 | 0.01 |
| V296 | 1.22 | 0.04 | 9.88 | 0.16 | 0.46 | 0.02 |
| T297 | 1.36 | 0.08 | 11.99 | 0.31 | 0.66 | 0.01 |
| I298 | 1.38 | 0.04 | 12.12 | 0.24 | 0.73 | 0.02 |
| E299 | 1.45 | 0.05 | 12.90 | 0.15 | 0.67 | 0.01 |
| S300 | 1.38 | 0.09 | 12.96 | 0.24 | 0.76 | 0.01 |
| V301 | 1.38 | 0.09 | 11.77 | 0.21 | 0.60 | 0.02 |
| A302 | 1.47 | 0.05 | 13.48 | 0.52 | 0.78 | 0.03 |
| D303 | 1.38 | 0.03 | 13.67 | 0.48 | 0.79 | 0.01 |
| Y304 | 1.45 | 0.04 | 11.61 | 0.17 | 0.79 | 0.01 |
| F305 | 1.36 | 0.09 | 12.34 | 0.32 | 0.76 | 0.02 |
| K306 | 1.38 | 0.04 | 12.43 | 0.21 | 0.72 | 0.01 |
| Q307 | 1.49 | 0.01 | 13.03 | 0.12 | 0.70 | 0.01 |
| I308 | 1.38 | 0.04 | 13.50 | 0.28 | 0.72 | 0.02 |
| G309 | 1.38 | 0.05 | 14.47 | 0.49 | 0.72 | 0.01 |
| I310 | 1.25 | 0.05 | 12.66 | 0.28 | 0.61 | 0.01 |
| I311 | 1.41 | 0.05 | 15.80 | 0.44 | 0.73 | 0.03 |
| T313 | 1.33 | 0.04 | 12.74 | 0.34 | 0.78 | 0.01 |
| N314 | 1.27 | 0.08 | 11.58 | 0.21 | 0.70 | 0.01 |
| T317 | 1.32 | 0.02 | 13.14 | 0.61 | 0.61 | 0.01 |
| G318 | 1.40 | 0.05 | 11.34 | 0.28 | 0.68 | 0.01 |
| Q319 | 1.36 | 0.09 | 10.69 | 0.13 | 0.60 | 0.01 |
| M321 | 1.19 | 0.10 | 10.95 | 0.50 | 0.73 | 0.08 |
| I322 | 1.34 | 0.05 | 12.98 | 0.44 | 0.74 | 0.02 |
| L324 | 1.32 | 0.07 | 11.39 | 0.31 | 0.70 | 0.02 |
| T326 | 1.29 | 0.04 | 13.53 | 0.32 | 0.71 | 0.02 |
| D327 | 1.32 | 0.05 | 13.92 | 0.15 | 0.72 | 0.01 |
| E329 | 1.37 | 0.08 | 13.66 | 0.13 | 0.53 | 0.005 |
| T330 | 1.26 | 0.02 | 28.96 | 0.82 | 0.54 | 0.01 |
| G331 | 1.44 | 0.05 | 9.53 | 0.32 | 0.26 | 0.00 |
| K332 | 1.40 | 0.05 | 4.40 | 0.07 | 0.08 | 0.00 |
| G335 | 1.37 | 0.08 | 19.73 | 1.05 | 0.72 | 0.03 |
| T338 | 1.40 | 0.06 | 15.21 | 0.82 | 0.80 | 0.01 |
| V339 | 1.40 | 0.04 | 11.80 | 0.40 | 0.77 | 0.04 |
| S340 | 1.41 | 0.08 | 11.96 | 0.16 | 0.74 | 0.02 |
| F341 | 1.41 | 0.05 | 12.70 | 0.48 | 0.79 | 0.02 |
| D343 | 1.41 | 0.07 | 7.70 | 0.37 | 0.15 | 0.00 |
| S346 | 1.44 | 0.03 | 12.24 | 0.08 | 0.77 | 0.01 |
| A347 | 1.42 | 0.05 | 14.28 | 0.43 | 0.76 | 0.04 |
| A349 | 1.47 | 0.04 | 5.39 | 0.56 | 0.82 | 0.03 |
| A350 | 1.44 | 0.06 | 13.95 | 1.00 | 0.81 | 0.03 |
| I351 | 1.38 | 0.08 | 13.15 | 0.31 | 0.80 | 0.02 |
| D352 | 1.39 | 0.07 | 14.16 | 0.26 | 0.76 | 0.01 |
| W353 | 1.37 | 0.08 | 12.16 | 0.17 | 0.78 | 0.01 |
| F354 | 1.38 | 0.07 | 13.89 | 0.19 | 0.78 | 0.02 |
| D355 | 1.39 | 0.08 | 13.99 | 0.15 | 0.76 | 0.01 |
| E358 | 1.41 | 0.08 | 5.68 | 0.16 | 0.30 | 0.00 |
| F359 | 1.32 | 0.05 | 15.99 | 0.46 | 0.75 | 0.02 |
| S360 | 1.38 | 0.07 | 15.23 | 0.67 | 0.75 | 0.02 |
| G361 | 1.33 | 0.02 | 14.22 | 0.18 | 0.68 | 0.01 |
| N362 | 1.37 | 0.06 | 11.54 | 0.22 | 0.71 | 0.01 |
| I364 | 1.58 | 0.09 | 12.40 | 0.14 | 0.66 | 0.01 |
| V366 | 1.41 | 0.06 | 11.96 | 0.60 | 0.75 | 0.04 |
| S367 | 1.37 | 0.08 | 11.90 | 0.10 | 0.78 | 0.01 |
| A369 | 1.44 | 0.09 | 11.11 | 0.27 | 0.76 | 0.03 |
| T370 | 1.26 | 0.09 | 10.25 | 0.08 | 0.62 | 0.01 |
| R372 | 1.50 | 0.07 | 7.56 | 0.34 | 0.27 | 0.01 |
| D374 | 1.53 | 0.05 | 5.97 | 0.09 | 0.23 | 0.003 |
